# Chronic warming and dry soils limit carbon uptake and growth despite a longer growing season in beech and oak

**DOI:** 10.1101/2023.07.18.549347

**Authors:** Margaux Didion-Gency, Yann Vitasse, Nina Buchmann, Arthur Gessler, Jonas Gisler, Marcus Schaub, Charlotte Grossiord

## Abstract

Progressively warmer and drier conditions impact tree phenology and carbon cycling with large consequences for forest carbon balance. However, it remains unclear how individual impacts of warming and drier soils differ from their combined one and how species interactions modulate tree responses. Using mesocosms, we assessed the multi-year impact of continuous air warming and lower soil moisture acting alone or combined on phenology, leaf-level photosynthesis, non-structural carbohydrate concentrations, and aboveground growth of young European beech and Downy oak trees. We further tested how species interactions (monocultures *vs*. mixtures) modulated these effects. Warming prolonged the growing season of both species but reduced growth for oak. In contrast, lower moisture did not impact phenology but reduced trees’ assimilation and growth for both species. Combined impacts of warming and drier soils did not differ from single ones. Performances of both species in the mixtures were enhanced compared to the monocultures under extreme conditions. Our work revealed that higher temperature and lower soil moisture have contrasting impacts on phenology *vs*. leaf-level assimilation and growth, with the former being driven by temperature and the latter by moisture. Furthermore, we show a compensation of the negative impacts of extreme events by tree species interactions.

## Introduction

With chronically rising air temperature, forests will be more frequently exposed to a combination of hotter and drier conditions in the future (Breshears et al., 2005), inducing a shift of the climatic range closer to the current distribution limit of species (i.e., rear edge). The co-occurrence of chronic warming and lower soil moisture could have contrasting impacts on climate-vegetation predictions as temperature and soil moisture can have different effects on forest carbon cycling in terms of phenology, photosynthesis and growth (Fu et al., 2020; Didion-Gency et al., 2022; Petrik et al., 2022; Vitasse et al., 2022). Hence, there is an urgent need to disentangle single from combined impacts of chronically elevated temperatures and lower soil moisture on tree carbon relations.

Most studies demonstrated that higher spring temperatures induce earlier leaf-out of trees potentially leading to a longer growing season (Polgar and Primack, 2011). However, the underlying leaf-level cues driving these effects and inter-specific differences are not fully understood (Schaber and Badeck, 2003). For instance, the winter bud dormancy of European beech (*Fagus sylvatica* L.) is sensitive to photoperiod and requires a high amount of winter chilling to be released (Way and Montgomery, 2015). On the contrary, some species, such as sessile oak (*Quercus petraea* (Matt.) Liebl) or oriental oak (*Quercus variabilis* Blume) are mainly sensitive to variations in spring temperature, showing large advancement of leaf emergence with increasing temperature (Dantec et al., 2014; Han et al., 2014). One of the simplest models to predict budburst timing is to compute the amount of heat necessary to trigger this phenological stage, classically expressed as growing degree day (GDD) above a certain temperature threshold (generally set at 5°C). However, previous work suggested that the amount of GDD reached at budburst varies depending on the previous exposure to chilling temperatures responsible for dormancy release in winter, i.e., exposure to temperatures ranging between *ca.* - 2°C to 10°C (Baumgarten et al., 2021). Thus, the GDD requirement gradually decreases with longer exposition to chilling conditions, reaching a minimum when chilling has been sufficient to fully release dormancy (Murray et al., 1989; Baumgarten et al., 2021). The interaction between chilling and heat requirement could explain the decline in phenological sensitivity to temperature increase recently found in European trees (Zhang et al., 2021), although this effect varies among species and may also be related to the non-linear effect of temperature (Wolkovich et al., 2021).

In contrast to spring phenology, the drivers of senescence timing are less understood. Contrasting patterns have been observed, including a warming-induced delay, advancement, or no effect on the date but an overall slower senescence process (Polgar and Primack, 2011). In addition, temperature impacts on phenology are not independent of soil moisture. For instance, low soil moisture can delay leaf emergence and cause premature senescence, thereby shortening the growing season (Bigler and Vitasse, 2021; Dallstream and Piper, 2021). A delay in leaf flushing has already been observed for beech trees under limited soil moisture and can be associated with a reduction of late frost risk and higher drought resistance by postponing the onset of transpiration (Spieß et al., 2012). Besides, a recent study suggest that the initiation of leaf senescence could largely be regulated by growth and development during early summer, whereas the leaf coloration and senescence progress is mediated by late summer/early autumn temperature (Zohner et al., 2023). Thus, the role of spring, autumn, and winter temperatures and their interactive effects with soil water availability remains unclear and needs to be explored further to better predict phenological trends.

Phenological shifts induced by warmer and drier soils affect the timing of leaf-level photosynthetic activity, tree growth, and carbon stocks in several ways (Polgar and Primack, 2011; Klein et al., 2016). For instance, an earlier leaf-out and extended growing season do not always result in an increase in carbon uptake and higher non-structural carbohydrate (NSC) concentrations at the yearly-scale because of an earlier start of assimilation (Etzold et al., 2022; Grossiord et al., 2022). Neither does it necessarily result in higher growth rates (Dow et al., 2022). Indeed, higher temperatures can harm trees when the optimal temperature for photosynthetic activity (T_opt_) is exceeded, resulting in reduced assimilation (Dai et al., 2021). Therefore, variation in these responses depends on the species-specific sensitivity to temperature. For instance, beech typically grows in a mild climate and has a T_opt_ around 24.5°C (Holišová et al., 2013), while downy oak (*Quercus pubescens* Willd.) usually grows in Mediterranean areas with a T_opt_ of about 26°C (Ilnitsky et al., 2021). Consequently, hotter and drier conditions can lead to severe changes in assimilation during the growing season (Etzold et al., 2022), which may be reflected in carbon stocks, including a reduction in NSCs pools (Hartmann and Trumbore, 2016) and growth (Morin et al., 2009). Moreover, warmer temperature co-occurs with a rise in vapor pressure deficit (VPD) and reduced soil water availability, which further restricts stomatal conductance (Zhou et al., 2014), resulting in lower growth, assimilation (Grossiord et al., 2020; Trotsiuk et al., 2021) and potentially depletion of NSC pools (Klein, 2015).

Yet, while the single impacts of lower soil moisture and warming have been studied in various ecosystems, the combined effects of chronically higher temperature and drier soils on tree phenology and carbon relations remain uncertain. Work conducted in drylands reported a delayed and prolonged bud development under combined hot and dry conditions associated with a more rapid depletion of NSCs (Adams et al., 2015). However, others have suggested an earlier spring phenology during hot droughts because of a more decisive temperature impact on phenology than soil moisture (Arzac et al., 2021). Although it remains untested, the contrast between these studies could be associated with species-specific tolerance to low soil water availability, the intensity of warming and soil moisture reduction, and the duration of these extreme conditions. Similarly, distinct responses have been observed regarding the impact of hot and dry conditions on leaf-level carbon relations. While most studies tend to agree that drier conditions have a more adverse impact on assimilation, NSCs, and growth (Lukasová et al., 2020), some studies have reported an exacerbation of trees’ responses under additive conditions (Arend et al., 2016). Thus, a better understanding of the combined impacts of chronic temperature rise and dry soils requires studies where single *vs*. additive warming and low moisture impacts on different species with contrasting strategies are compared over multiple years.

Although better predicting the fine-scale mechanisms driving tree responses to continuous warmer and drier conditions is fundamental, these responses cannot be fully understood without considering the dynamics of the forest as an entire system, particularly when considering species diversity (Forrester, 2014). Numerous studies have shown that forests with more than one tree species (i.e., mixtures) are often more resistant to extreme conditions than single-species forests (i.e., monocultures) (Vacek et al., 2021). Although these observations are species- and context-specific, they suggest that adverse impacts from chronic heat and low moisture could be mitigated in diverse forests. Two underlying mechanisms explaining biodiversity effects are usually applied. The “complementarity effect” is associated with species’ niche differences and/or facilitation. For instance, complementarity occurs when trees with distinct crown architectures and light requirements, such as beech and downy oak, interact, leading to a more efficient above-ground space occupation and increased biomass (Jucker et al., 2015). The “selection effect” reflects the dominance of more competitive species in mixtures (Loreau and Hector, 2001; Grossiord et al., 2013), resulting, for instance, in a lower reduction of productivity in mixtures compared to monocultures during drought (Dziedek et al., 2016). An approach to estimate the relative contribution of these two processes is to separate the net biodiversity effects into complementarity and selection effects (Loreau and Hector, 2001). However, to date, no studies investigated if and how tree species interactions affect phenological, leaf-level carbon cycle, and tree growth responses to warming and low soil moisture acting individually or combined.

Here, we aim to understand how carbon relations, including phenology, leaf-level gas exchange, and growth traits, of two co-occurring widely distributed and contrasting tree species, i.e., European beech and downy oak, are impacted by chronic air warming and a moderate but continuous reduction of soil moisture acting alone or combined over multiple years, and how tree species interactions can alter these responses. We exposed three-year-old beech and downy oak trees in monocultures and mixtures to a continuous +5°C air warming and a reduction of soil moisture by 50%, acting individually or combined using open-top chambers for three years. European beech and downy oak trees were selected because they can be found growing together in natural ecosystems, have different phenological cycles (Baumgarten et al., 2021), and different strategies to deal with low soil moisture and high temperature, with downy oak being more tolerant to moisture limitation and heat (Barigah et al., 2013).

Our objectives were to (1) evaluate how beech and downy oak phenology (including bud swelling, duration of bud development, onset and duration of senescence, growing season length) respond to chronic warming and moderate reduction of soil moisture acting individually or combined, (2) assess how these climatic conditions influence leaf-level carbon relations (i.e., starch and sugar concentrations, light-saturated assimilation, A_sat_) and growth (height, diameter, and aboveground biomass (AGB) increment), and finally (3) determine if and through which mechanisms species interactions (i.e., monocultures *vs.* mixtures) influence chronic warming and reduction of soil moisture impacts on phenology, leaf-level carbon relations, and growth. Because of the strong photoperiodic control of spring phenology in beech compared to downy oak (Basler and Körner, 2012), we expect (1) beech trees to show a weaker phenological shift in response to warming compared to downy oak trees, manifested through earlier leaf-out and slower senescence process inducing a longer growing season. On the contrary, we expect reduction of soil moisture to delay leaf development and advance leaf senescence, especially for the more moisture-sensitive beech trees. Combined climatic treatments on phenology should differ between these two species as temperature changes more strongly drive downy oak. Hence, a weaker advance in leaf-out and senescence are expected for beech, while a faster leaf emergence and slower senescence process could be observed in oak. Moreover, because of beech’s sensitivity to high temperature and low soil moisture, we expect (2) beech to have impaired assimilation, lower starch and sugar concentrations, and aboveground growth under chronic warming and reduced soil moisture. On the other hand, while drier soils should reduce these traits in downy oak, chronic warming may enhance them because of its higher tolerance to temperature reflected in its distribution range. Combined treatments should exacerbate tree responses observed under drier soils because of enhanced water stress. Finally, we expect (3) both species to benefit from being grown in mixtures compared to monocultures under all climatic conditions because of complementarity in their crown architectures and light requirements.

## Results

### Phenology

We found substantial impacts of chronic warming but not soil moisture on both species’ phenology (Fig. **1**, Fig. **S1**). Higher daily air temperatures in spring were linked to an earlier bud swelling for both species, where beech was advanced by −1.1 days.°C^-1^ and oak by −2.3 days.°C^-1^ across the two years and treatments. A shorter leaf development duration was also found, with a reduction of −0.7 days.°C^-1^ for both species. Similarly, senescence duration was extended by +3.4 days.°C^-1^ for beech and +3.1 days.°C^-1^ for oak. The growing seasons were extended by +4.5 days.°C^-1^ and +5.7 days.°C^-1^ for beech and oak, respectively, with an increase in daily mean annual air temperature (Fig. **1**). However, oak trees were taking more advantage from chronic warming compared to beech, as the advancement of bud swelling extension was higher (p = 0.004 using Least-squares analysis). No relationship between the onset of senescence and the fall daily air temperatures was found, suggesting that the onset of senescence is not driven by temperature changes in the fall for both species. We observed no significant relationship between phenology and soil moisture for both species (Fig. **S1**), indicating that temperature has a more direct impact on tree phenology than soil moisture.

**Figure 1:**
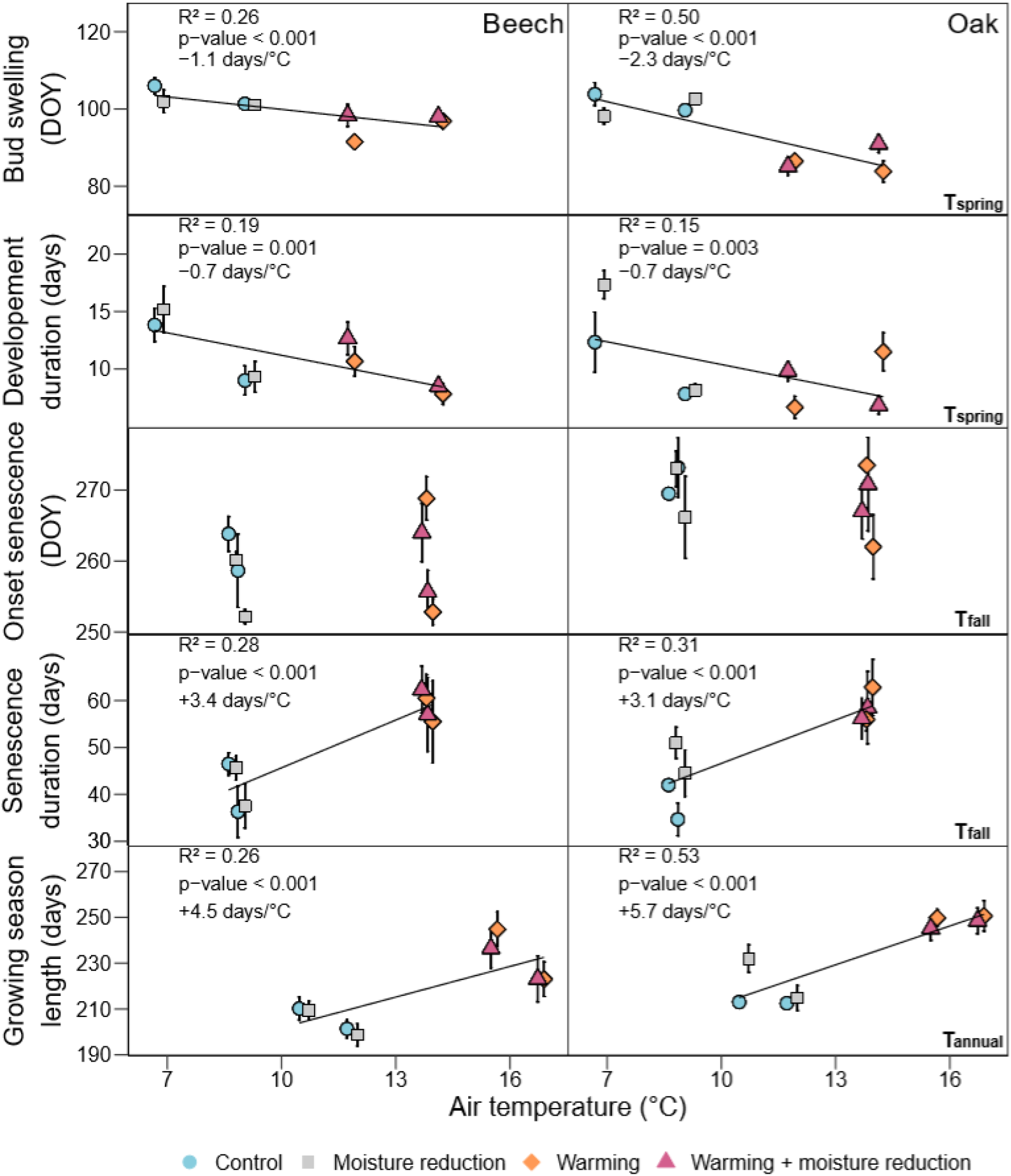
Bud swelling, leaf development duration, onset of senescence, leaf senescence duration, and growing season length as a function of air temperature for beech and oak trees growing under control (blue), moisture reduction (grey), warming (orange), and warming + moisture reduction (purple) in monocultures in 2020 and 2021 (mean ± SE per treatment, species and years, n = 6 trees). The bud swelling and leaf development duration are shown as a function of the daily mean air temperature in spring (from February to April, T_spring_). Onset of senescence and leaf senescence duration are shown as a function of the daily mean air temperature in fall (from September to December, T_fall_). Growing season length is shown as a function of the daily mean annual air temperature (T_annual_). Linear regression lines across all treatments per species are shown when significant. R², p-values and the change of day number for each degree are given in the top left corner when significant.

With an increase in the number of chilling days, both species had lower GDD requirements to budburst (Fig. **2**), suggesting that trees exposed to chronic warmer winters in the warm and low soil moisture conditions were likely limited in chilling to fully break dormancy (i.e., to have the minimum of forcing requirement to budburst).

**Figure 2:**
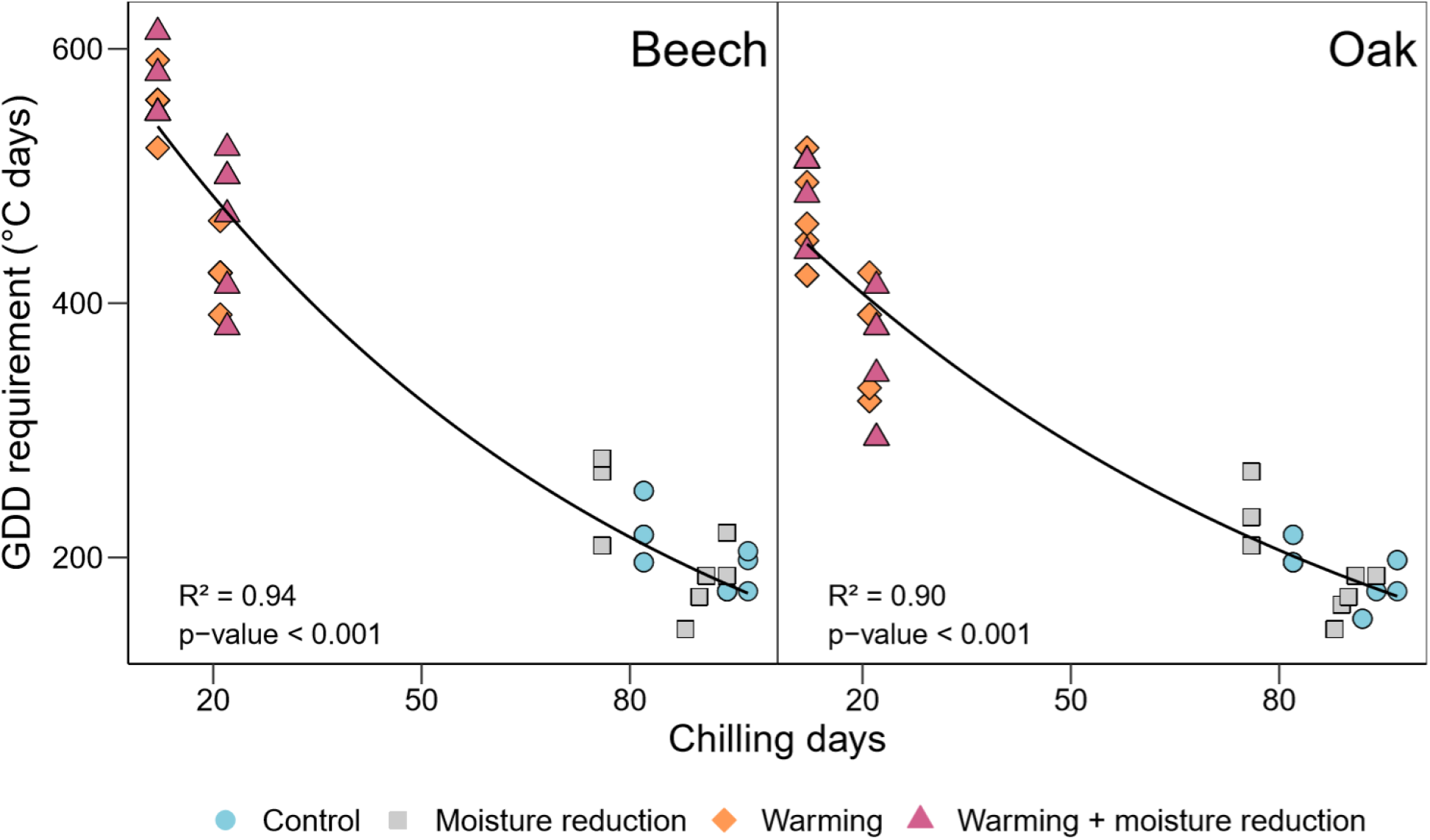
Growing degree day (GDD) requirement (i.e., accumulated the daily mean temperature above a threshold of 5°C from January 1^st^ to the date of bud swelling) as a function of the number of chilling days (i.e., days with average temperature below 5°C from the November 1^st^ to the date of leaf flushing) for beech and oak trees growing under control (blue), moisture reduction (grey), warming (orange), and warming + moisture reduction (purple) in monocultures in 2020 and 2021 (each point represents one tree). Negative exponential regression lines across all treatments per species are shown when significant. R² and p-values are given in the bottom left corner when significant.

### Leaf water potential

We found little effect of warming and soil moisture acting alone on predawn and midday leaf water potentials (Ψ_PD_ and Ψ_MD_, respectively, Fig **3**). Warming resulted in more negative Ψ_PD_ in 2020 for both species, as well as more negative Ψ_MD_ in 2020 but for beech only (Fig. **3**). Moreover, lower soil moisture reduced Ψ_PD_ in 2020 but for oak only (Fig. **3**), suggesting that the reduction in soil moisture was moderate. However, combined warming and drier soils resulted in more negative Ψ_PD_ in 2021 for both species (Fig. **3**, Table **S1**).

**Figure 3:**
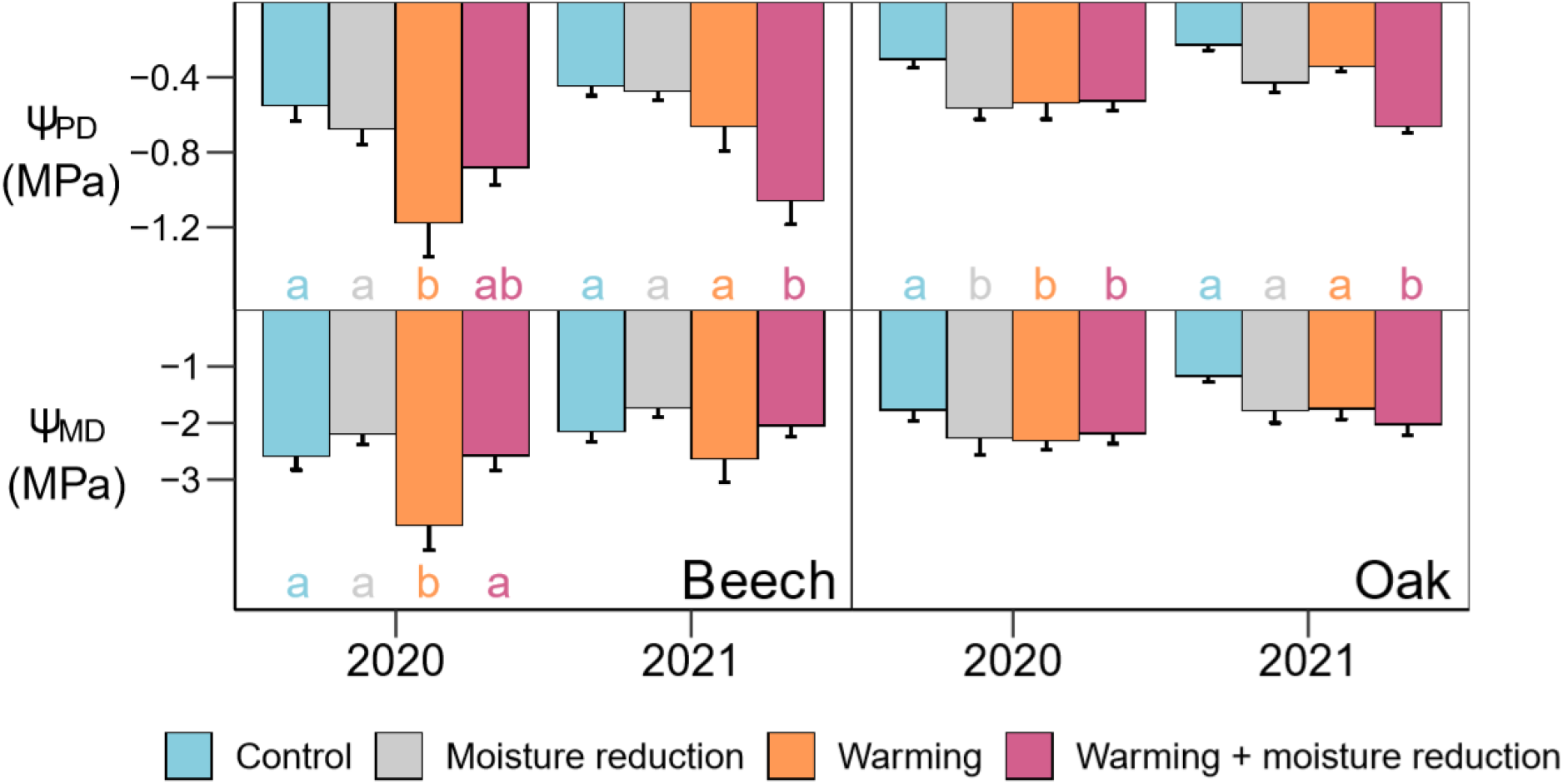
Predawn (Ψ_PD_) and midday leaf water potential (Ψ_MD_) for beech and oak trees growing under control (C, blue), moisture reduction (MR, grey), warming (W, orange), and warming + moisture reduction (WMR, purple) in monocultures in 2021 (mean ± SE per treatment and species, n = 6 trees per histogram). Significant differences between climatic treatments are highlighted for each species with letters (Tukey’s HSD post-hoc test, alpha = 0.05).

### Leaf-level carbon relations and tree growth

While lower soil moisture reduced level-level assimilation and growth, we found only a few impacts of chronic warming (Fig. **4**). For beech, warming had no impact on leaf-level carbon relations and growth (Fig. **4**, Table **S1**). For oak, warming reduced height increment, diameter increment, and aboveground biomass (AGB) increment but in 2021 only (Fig. **4**, Table **S1**). On the contrary, for beech, lower soil moisture led to a significant reduction of A_sat_ in 2021, height increment in 2021, diameter increment in 2020 and 2021, and AGB increment in 2021. Similarly, for oak, we observed a substantial reduction in A_sat_, height increment, diameter increment, and AGB increments in 2020 and 2021 (Fig. **4**, Table **S1**). Under combined warming and low moisture, we observed a reduction of A_sat_, and diameter increment in 2020 and 2021 for beech. For oak, a reduction of A_sat_, height increment, diameter increment, and AGB increment in 2020 and 2021 was observed (Fig. **4**, Table **S1**). No significant differences were found between lower moisture alone and combined warming and soil moisture reduction responses (Fig. **4**, Table **S1**).

**Figure 4:**
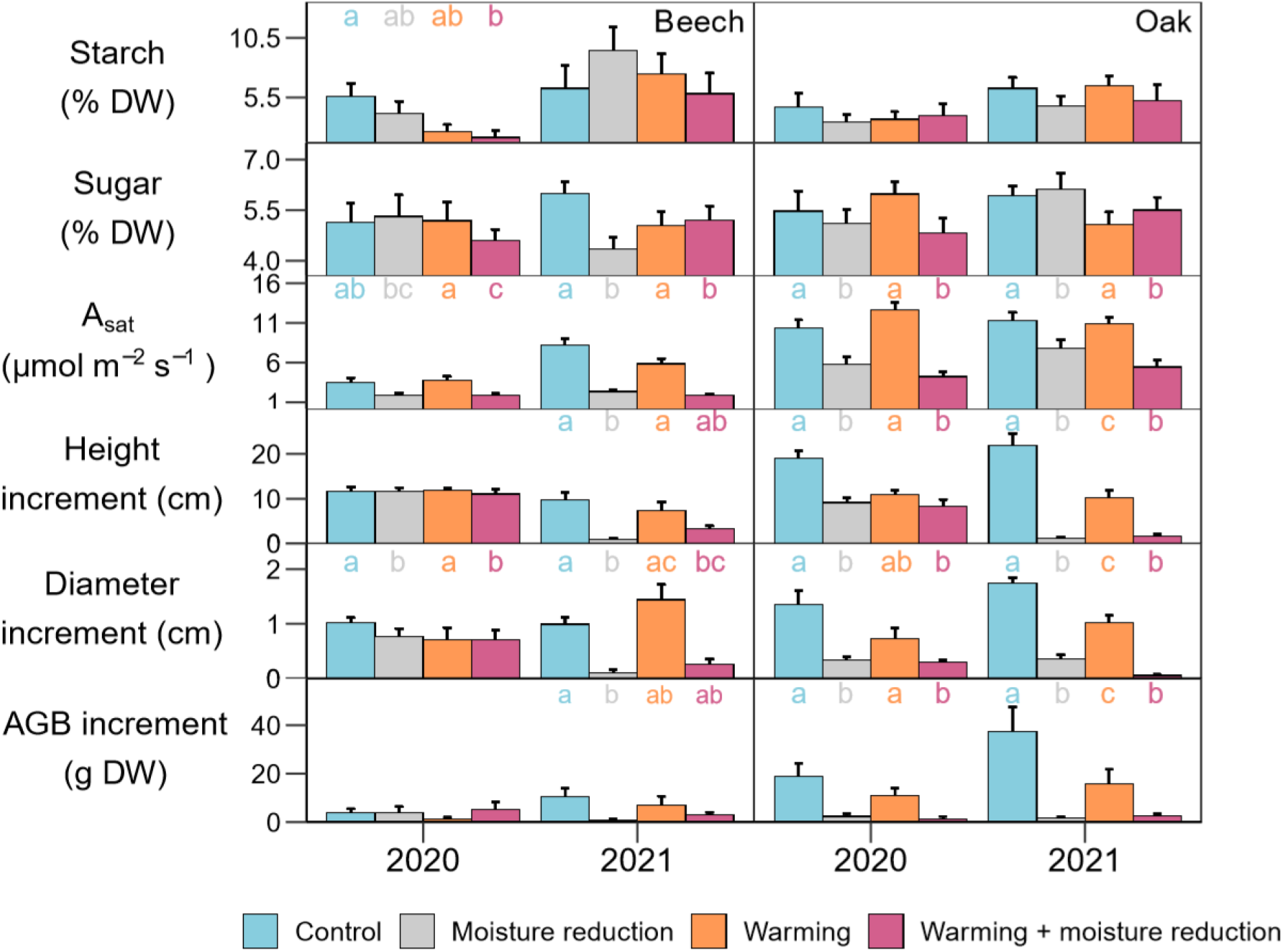
Starch, sugar, light-saturated assimilation (A_sat_), height increment, diameter increment, and estimated aboveground biomass (AGB) increment for beech and oak trees growing under control (C, blue), moisture reduction (MR, grey), warming (W, orange), and warming + moisture reduction (WMR, purple) in monocultures in 2021 (mean ± SE per treatment and species, n = 18 trees per histogram for the physiological traits and n = 6 trees per histogram for the growth traits). Significant differences between climatic treatments are highlighted for each species with letters (Tukey’s HSD post-hoc test, alpha = 0.05).

### Impact of species interactions

For both species, no impact of species interactions on phenology was found in all treatments, except for oak where we observed a slightly shorter development duration of 2 days in mixtures compared to the monocultures under lower moisture (Table **S2)** but no significant net biodiversity effect. Similarly, we observed limited species interaction effects in control or when warming and lower soil moisture acted alone in 2020 and 2021. However, a positive effect of species mixture under combined warming and drier soils was found (Fig. **5**).

**Figure 5:**
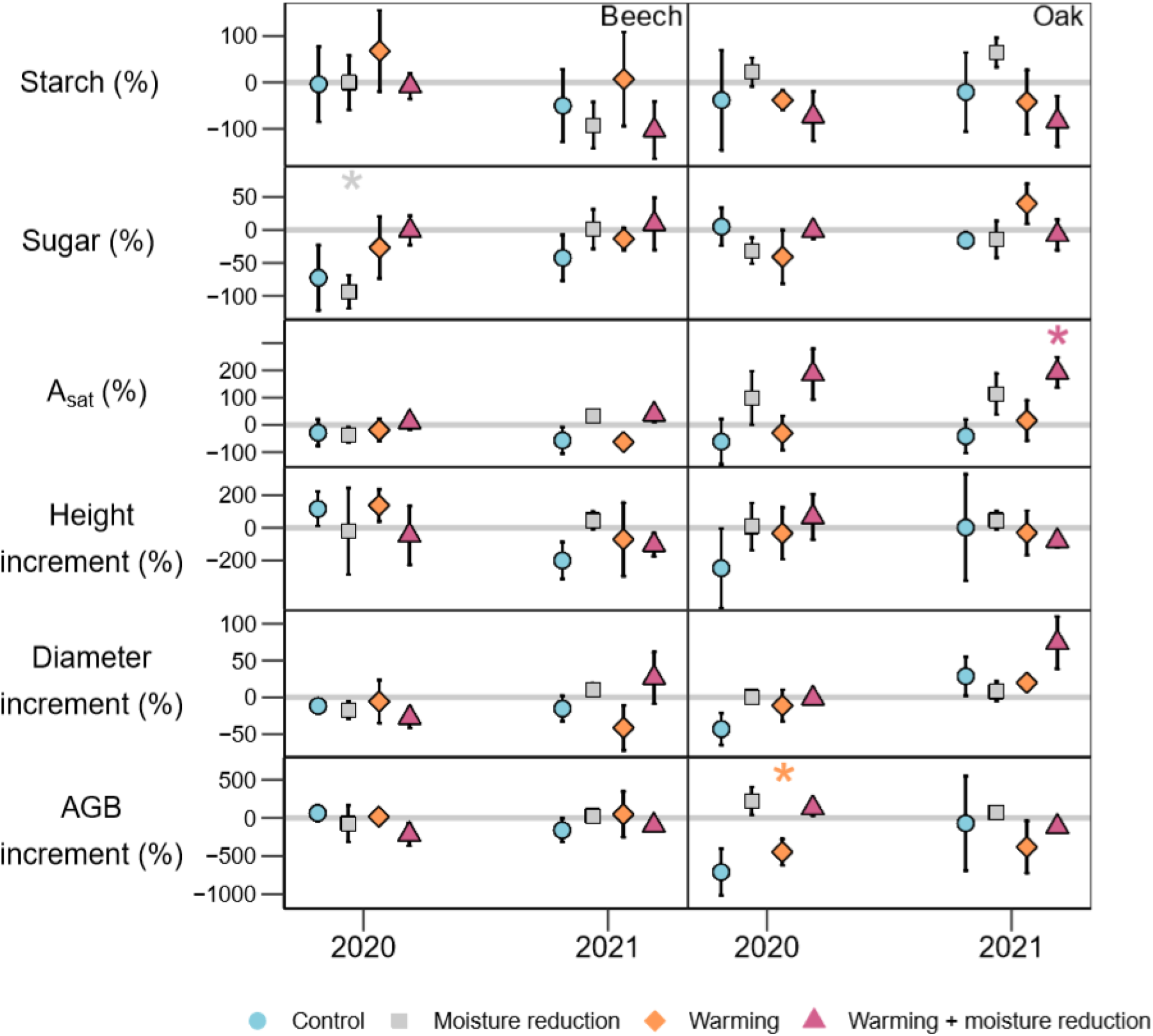
Starch, sugar, light-saturated assimilation (A_sat_), height increment, diameter increment, and estimated aboveground biomass (AGB) increment net biodiversity effect for beech and oak trees growing under control (C, blue), moisture reduction (MR, grey), warming (W, orange), and warming + moisture reduction (WMR, purple) (mean ± SE per treatment and species for the years 2021, n = 18 trees per symbol for the physiological traits and n = 6 trees per symbol for the growth traits). Positive values indicate higher rates in mixtures compared to the monocultures. Significant differences from 0 are highlighted per treatment and species (p - value: 0.05 ≥ * > 0.01, 0.01 ≥ ** > 0.001, *** ≥ 0.001).

For beech, no impact of species interactions on leaf-level carbon relations, or growth was found in the warming treatment in 2020 and 2021 (Table **S3**). Consequently, beech trees did not show any impact of net biodiversity in response to warming in terms of leaf-level carbon relations or growth in 2021 (Fig. **5**). However, we observed a positive selection effect on beech A_sat_ under warming (Fig. **S2**), suggesting positive interactions in the mixtures. Although this effect was not strong enough to influence the overall net biodiversity effect. For oak, no differences between mixtures and monocultures were found under warming, except for lower diameter increment of about 0.3 cm per year in mixtures compared to monocultures (Table **S3**). Moreover, oak trees showed a negative net biodiversity effect for the AGB increment in response to warming in 2020 (Fig. **5**), explained by a negative complementarity effect (Fig. **S3**).

Under lower soil moisture, we found a reduced diameter increment in mixtures compared to the monocultures for beech (Table **S3**). Similarly, a negative net biodiversity effect on sugar concentration in response to low soil moisture was found on beech (Fig. **5**), driven by a negative complementarity effect (Fig. **S3**). In contrast, we observed no impact of species interactions and net biodiversity effect on oak (Table **S3**, Fig. **5**).

Under combined warming and lower moisture, a larger height increment by about 2 cm per year was found in monocultures compared to the mixtures for beech (Table **S3**). On the contrary, for oak, diameter increment was larger by 1.4 cm in mixtures compared to the monocultures (Table **S3**). We also observed a positive net biodiversity effect for A_sat_ in 2021 for oak (Fig. **5**). This effect was explained by complementarity effects, indicating a resource complementarity under warmer air and drier soils (Fig. **S3**).

## Discussion

Contrary to our expectations, chronic warming had similar effects on the spring phenology advancement of both species. Bud swelling was advanced by around −1.1 and −2.3 days.°C^-1^ for beech and downy oak, respectively (Fig. **1**), confirming previous work for beech (Dantec et al., 2014). These new findings for downy oak suggest a similar response to continuous warming as other previously studied oak species (e.g., sessile oak - Charlet de Sauvage et al., 2022; and oriental oak - Han et al., 2014). Budburst onset in beech was less sensitive to temperature than in downy oak, which was assumed because of the stronger photoperiod and chilling temperature dependency compared to oak species (e.g., common oak (*Quercus robur* L.) and sessile oak - Lebourgeois et al., 2010; Fu et al., 2013). Moreover, we found that a lower amount of winter chilling increases the demand for bud flushing for cumulative forcing temperatures in spring (Fig. **2**), as previously assumed (Murray et al., 1989). However, while most studies showed an exponential relationship between GDD and chilling days, suggesting that trees exposed to a lower number of chilling days require higher temperatures in spring to flush, our results indicate that the climatic conditions of the open-top chambers might be in the more linear part of the negative exponential relationship between these two parameters. This result suggests that the winter temperature did not fulfill the required chilling for both species. A study showed that beech and sessile oak need more than 160 and 90 days (Vitasse and Basler, 2013), respectively, with a temperature below 5°C to reach the chilling requirement. In our experiment, trees were exposed to a maximum of 97 days even in the absence of warming (Fig. **2**), which could partially explain the more extensive heat-induced budburst advancement in downy oak compared to beech, as another oak species, i.e., sessile oak, has already shown lower chilling requirements than beech. Nevertheless, while trees may not have been ultimately released from winter dormancy, higher spring temperatures in the warmed treatments compensated for the warming-induced reduction in chilling and still induced an earlier bud break in both species, as already found (Zhang et al., 2021). However, these findings should be interpreted with caution because photoperiod is another parameter that affects the relationship between chilling and GDD requirements (Fu et al., 2019). While further experiments with distinct gradual chilling and forcing temperatures would be needed to better disentangle the environmental cues driving budburst, our results suggest that warmer springs alone significantly promote the start of the growing season in tree species with contrasting temperature and moisture tolerances. Hence, we expect the trend in spring advancement observed over past decades (Vitasse et al., 2018) to continue with the ongoing shift towards warmer conditions, irrespective of higher winter temperatures, but likely at a slower rate due to chilling and photoperiodic limitations (Fu et al., 2015) or due to non-linear effects of temperature increase on development (Wolkovich et al., 2021).

Moreover, while our treatments did not change the onset of senescence, we found that the process of senescence and the growing season was extended for both species with warming (Fig. **1**). Previous work reported similar results (Jiang et al., 2022). Leaf senescence is driven by the interaction of multiple environmental factors, including light conditions (Vitasse et al., 2021), cold and warm temperatures (Schuster et al., 2014), and soil moisture (Holland et al., 2016), with some species being more sensitive to specific parameters than others. Our results indicate that beech and oak trees’ senescence show minor sensitivity to chronic temperature changes and may, therefore, be more dependent on the photoperiod. Nevertheless, according to a recent work from Zohner et al. (2023), pre-summer solstice warming accelerates senescence, while post-solstice warming delays it. Thus our all year long warming applied could have compensate these opposing effects. In consequence, it is essential to note that our experiment did not allow us to distinguish the single effects of spring, fall, and winter temperatures because warming was applied throughout the experimental period to reflect changes in the climatic range towards warmer and drier conditions. Further studies applying warming only in spring, fall, and/or winter would allow us to better understand the exact cues driving senescence and/or the impact of single heat waves to incorporate phenological shifts in models that do not consider climate-induced phenological changes.

Contrary to temperature, we found no effect of lower soil moisture on the phenology of both species (Fig. **S1**). We anticipated a shorter growing season under drier soils, as previously described in several studies (Adams et al., 2015), also including oaks species (Spiess et al., 2012; Dallstream and Piper, 2021). This finding suggests that water availability effects on phenology vary depending on the moisture level. For instance, using an experimental approach, one study showed that a 60% soil water reduction in acidic and calcareous soils induced a 2 days earlier leaf flushing in different oak provenances (i.e., sessile, common, and downy oak; Kuster et al., 2014). On the contrary, in a dry Mediterranean forest, others found that 30% precipitation reduction delayed budburst in holm oak (*Quercus ilex* L.; Limousin et al., 2012). As the predawn (Ψ_PD_) and midday leaf water potential (Ψ_MD_) of our trees were slightly affected by our low moisture treatment (Fig. **3**), we consider the water availability level as moderate. Indeed, trees reached Ψ_PD_ of about −1.9 and −1.2 MPa for beech and downy oak, respectively. In comparison, in a temperate forest in Switzerland, researchers found Ψ_PD_ values between −2 to −3.3 MPa for adult beech trees during the 2018 drought, which led to premature leaf senescence in late July (Schuldt et al., 2020). Hence, our work suggests that exposure to moderate soil moisture conditions has limited impacts compared to extreme dryness and that moisture impacts on phenology do not seem to follow any relationship. Moreover, similar responses were found under warmer and drier conditions (Fig. **1**), supporting our finding that a chronic +5°C warming has a more decisive impact on phenology than moderate reduction in soil moisture.

Warming under well-watered conditions had no effects on the leaf-level carbon relations and growth of beech (Fig. **4**). This finding contradicts our expectations and previous studies suggesting high heat vulnerability in beech (Gessler et al., 2006). Trees experienced a yearly MAT of maximum of 17°C (in 2020) in the warmed treatments, which is significantly above their optimum temperature range. However, beech can occur in environments where MAT reaches a maximum of 18°C (Durrant et al., 2016). Hence, our constant treatments may not have been strong enough to induce significant shift to this species’ rear edge. However, contrary to our expectations, downy oak, which occurs in environments of 19°C MAT (Pasta et al., 2016), showed no effect of warming on A_sat_ and NSCs, but a substantial decrease in growth (Fig. **4**). While this result contradicts previous studies where oak increased A_sat_ with a + 0.8°C and + 1.5°C warming (Arend et al., 2016), it supports recent studies finding reduced growth with higher temperature and VPD (Adams et al., 2015; Trotsiuk et al., 2021). However, growth reduction was not associated with lower carbon fluxes and storage (i.e., starch and sugar concentration, and A_sat_). While no respiration measurements were conducted in our study, we expect this trend to be partially driven by warming-enhanced leaf cellular respiration, as observed in previous studies (Piao et al., 2008). Moreover, temperature could increase soil respiration and thereby reduce the overall available carbon pools. Further research on whole-tree carbon losses through respiration, including the offsets in carbon gained during photosynthesis, would be necessary to assess chronic warming impacts on the carbon balance in this experiment.

On the contrary, chronically drier soils strongly reduced the leaf-level carbon relations and growth of both species, except for NSCs, where we found no changes (Fig. **4**). However, while both species were negatively impacted by low soil moisture, beech A_sat_ was more strongly impaired (Fig. **4**), which corresponds well with its rather isohydric stomatal behavior relative to others oak species (i.e., sessile, white and red oaks; Pretzsch et al., 2013). These findings support many studies showing that reduced soil moisture negatively affects A_sat_ and growth (Scharnweber et al., 2011), and indicate that although the reduction in soil moisture was moderate, it was sufficient to affect leaf carbon relations and growth of both species negatively. Homeostatic NSC concentrations under drier soils have also frequently been observed (Schönbeck et al., 2018) and could suggest that newly assimilated carbon was preferentially invested in storage and maintenance instead of growth. However, further measurements using, for instance, carbon isotopes would be needed the validate this assumption.

As observed for the phenological responses, no differences were observed for all physiological and growth traits under combined treatments and single stressors (Fig. **4**). However, contrary to phenology, which was driven mainly by temperature, leaf-level carbon relations and growth responded similarly as under drier soils only. Thus, our results indicate that water resources are a stronger determining factor for these responses than a continuous +5°C warming. These findings support previous work (Didion-Gency et al., 2021) and reinforce previous results where warming does not necessarily worsen soil moisture levels (Grossiord et al., 2017). Exacerbated impacts of warming imposed on dry soils have been associated with faster exhaustion of soil moisture through VPD-enhanced transpiration and higher residual water loss, accelerating dehydration after stomatal closure. We found no differences in soil moisture conditions at 25 cm depth between single and combined treatments (Fig. **6**), suggesting that both processes did not occur in our study. While this result could be explained by the relatively high relative humidity inside the chambers, the extreme dryness of the soil in the superficial soil layers probably prevented any further evaporative water loss. Moreover, residual water loss sensitivity to soil moisture has often been observed under drier conditions after a few years (Grossiord et al., 2018; Grossiord et al., 2020) and could be taken place at our site in the future. Plastic trait responses are often involved in climate change studies, but rarely to understand the responses to climatic range shifts toward the rear edge of a species. Our results further suggest that trees’ responses to chronic warming and low moisture depend on the stress duration as most effects were found after three years of treatment exposure (Fig. **4**). We show that physiological traits have a relatively slower adjustment compared to the more rapid phenological response. Indeed, a shift in the spring and autumn phenology was observed in the subsequent year following the initiation of the treatments (in 2020) while gas exchange and growth mainly deviated from the control after two years (in 2021). These findings support previous work showing shifts in phenology only one year after exposure to a new temperature regime (e.g., Morin et al., 2010). Similarly, studies that have artificially exposed trees to chronic mild drought over several years also found gas exchange and growth rates to mainly respond after two years (e.g., Schönbeck et al., 2020). Furthermore, it is essential to note that our work was conducted with trees at the seedling life stage and that adult trees could respond differently to warmer and drier conditions. For instance, it is important to notice that young beech and oak trees have marcescent leaves, which provide protection from freezing temperatures, but also reduce the exposure to heat, through a reduction of the temperature exchange between the stem and the surrounding environment, and potentially delay budburst timing, independently of the climatic conditions (Heberling and Muzika, 2023). Similarly, previous work found that young trees tend to have an earlier budburst as a compensatory mechanism for the overshadowing by taller trees in mature forests (Augspurger and Bartlett, 2003; Vitasse, 2013). However, studying juvenile phenology is still crucial for understanding trees establishment and predicting carbon sequestration in forests, as young trees are the key stage where rapid evolutionary responses occur.

**Figure 6:**
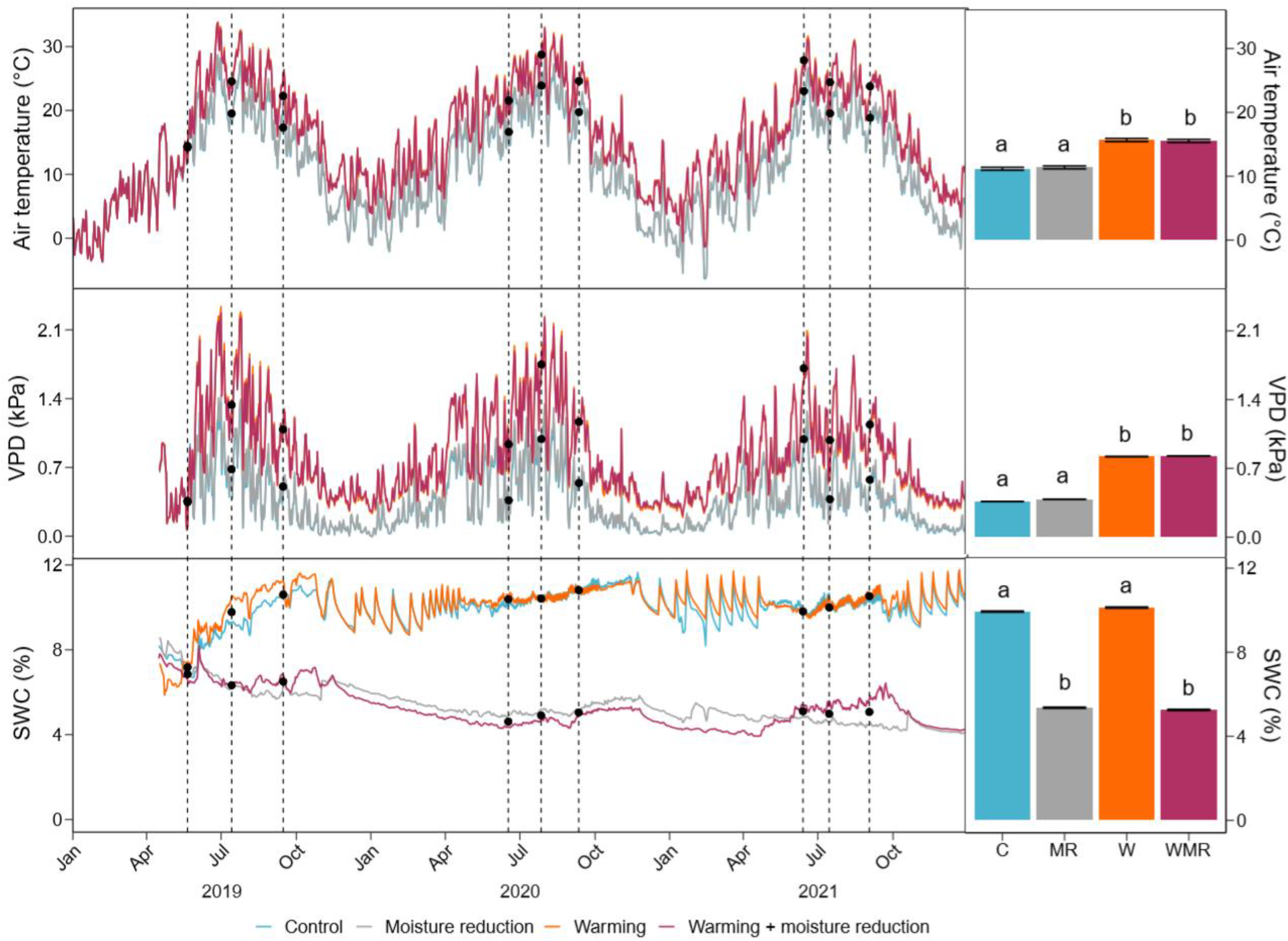
Air temperature, vapor pressure deficit (VPD), and soil water content (SWC) under control (blue), moisture reduction (grey), warming (orange), and warming + moisture reduction (purple) treatments measured in the open-top chambers. Full lines show the daily averages (n = 4 chambers per treatment). Dashed vertical lines indicate the middle of the measurement campaigns (lasting approx. 6 days). Black filled dots stand for the non-heated and heated chambers for the air temperature and VPD, and high soil moisture and low soil moisture chambers for the SWC during the campaigns (n = 8 each). Color-filled bars show the averages (mean ± SE) over the entire time in each climatic treatment, and the significant differences are highlighted with letters (Tukey’s HSD post-hoc test, alpha = 0.05).

We expected positive interactions between species because beech and downy oak trees exhibit contrasting functional strategies for resources uptake (e.g., water) and light use, leading to a possible reduction of competition for resources (Fabiani et al., 2022). Yet, species interactions did not impact phenology, suggesting that micro-climatic changes induced by biotic interactions were insufficient to impact the timing of flushing, senescence, and the growing season length (Table **S2**). However, depending on the climatic treatments and species, tree species interactions positively affected leaf-level carbon relations (Table **S3**).

When exposed to warming, beech showed no differences between mixtures and monocultures, and net biodiversity were close to zero for leaf-level carbon relations and growth traits (Table **S3**, Fig. **5**). On the contrary, oak suffered in mixtures relative to the monocultures in 2020 (Fig. **5**), which was driven by a positive complementarity effect (Fig. **5** & **S3**). This finding could suggest a higher competitive ability of beech at an early life stage. However, care must be taken as trees may still have been impacted by their transplantation in the chambers during the first year. Some oak species, such as northern red oak (*Quercus rubra* L.), are known to be highly sensitive to transplant shock because of to their taproot systems (Harris et al., 2002). Moreover, no impact was observed during subsequent years under warming. Under drier soils only, a slight shorter development duration in mixtures compared to the monocultures for oak trees, as well as a lower diameter increment in mixtures compared to the monocultures for beech were found (Table **S2** & **S3**), Similarly, a negative net biodiversity effect on sugar concentration in response to low water availability, driven by a positive complementarity effect was observed in 2020 (Fig. **5** & **S3**). Overall, these results suggest little impact of tree neighbors on leaf-level carbon relations and growth when drier soils act alone.

However, when warming and lower moisture co-occurred, oak trees had higher A_sat_ in mixtures compared to the monocultures after three years, leading to a positive net biodiversity effect (Fig. **5**), driven by complementarity between species (Fig. **S3**). These findings support previous work showing strong resource partitioning between beech and downy oak and beneficial interaction effects during extreme events (Grossiord et al., 2015). Nevertheless, the effect of species interactions was not constant throughout the experiment and treatment, indicating a shift in the type of interactions over time. Potentially, beech trees had faster root development after planting, allowing better access to soil nutrients and water resources in the first two years (2019 and 2020). One study observed similar results where more rapid root development was found in beech compared to oak when grown together at a young life stage (Leuschner et al., 2001). However, further work on root growth would be required to validate this hypothesis. Nevertheless, the positive complementarity effect found for oak after three years supports many studies suggesting the interactions between trees are changing over time, and often getting stronger (Domisch et al., 2015; Haase et al., 2015). Indeed, our experiment is still at an early stage of tree development (6 year-old trees), and trees interactions can take multiple years to establish (Domisch et al., 2015). Thus, further measurements would be needed to determine how the interactions will change with tree age and to confirm the apparent advantage of oak in older mixtures with beech during hotter and drier conditions.

Overall, our study highlights that European beech and downy oak will extend their growing season with chronically rising temperatures, even under lower soil moisture, with a longer extension for oak. Therefore, oak may become more competitive, especially in spring, as an earlier bud break and photosynthetic activity would provide access to resources before other species start their growing season. Moreover, while we found that a chronic +5°C warming will reduce the chilling conditions in winter required for the release of winter bud dormancy, a stronger forcing due to warmer springs will compensate for cold requirements in both species and spring phenology will thus contribute to advance bud break but at a slower rate as before. Contrary to phenology, which was not impacted by drier soils, prolonged water shortage will severely reduce tree gas exchange and growth with stronger impacts on beech. However, although temperature and low soil moisture impacted the phenology and leaf-level carbon relations differently, their additive effects did not differ from their single ones in our experiment. Therefore, our work suggests that trees could have lower carbon uptake during hotter and drier conditions and that an extension of their active period through chronic temperature rise could not compensate for this reduction as lower growth is still observed overall. Nevertheless, large uncertainties remain regarding the concurrent CO_2_ fertilizing impact on all these processes in the longer term, and future work should focus further on those additive climatic drivers (see Zani et al., 2020). Furthermore, we showed that species interactions could actively shape tree responses after a few years, with a positive effect of mixtures compared to monocultures, but only under extreme conditions.

## Methods

### Experimental set-up

The study was conducted in OTCs designed to investigate the impact of tree species interactions under chronic warming and low soil moisture (Didion-Gency et al., 2022). The site is located at the Swiss Federal Research Institute for Forest Snow and Landscape Research (WSL) in Birmensdorf, Switzerland (47°21’48” N, 8°27’23” E, 545 m a.s.l), and contains 16 hexagonal glass-walled OTCs (3 m height, 6 m² area each, Fig. **7**), where mobile roofs are kept above the chambers during the entire experiment to exclude natural precipitation and control the soil water status. The glass walls, roofs, and shadows between chambers reduce the photosynthetic active radiation (PAR) inside the OTCs by about 50% compared to the outside conditions (but still reach up to 1700 µmol m^-2^ s^-1^ PAR during sunny days). The belowground part of each OTC is divided into two semicircular lysimeters (1.5 m deep, 2.5 m²) with concrete walls, which are divided into 4 compartments using plexiglass walls, leading to a total of 8 soil compartments per OTC (each with an area of 0.625 m²). The soil compartments were filled with a 1 m deep layer of gravel for fast water drainage, covered with a fleece to prevent root growth into the gravel below the soil layers but to allow the water to pass through, and topped with 50 cm of an artificial forest soil provided by the company Ökohum (Herbertingen, Germany; pH 6.3, 40% quartz sand, 20% white peat, 20% expanded shale, 16% pumice stone, and 4% clay). In all compartments, annual soil fertilization was conducted in spring using granules (Unikorn I, Hauert, Grossaffoltern, Switzerland) with an amount of 30 g per compartment (20% potash, 14% nitrogen, 12% sulfur, 4% phosphate, 3% magnesium). During the entire experiment, the leaf litter was left in each compartment.

**Figure 7:**
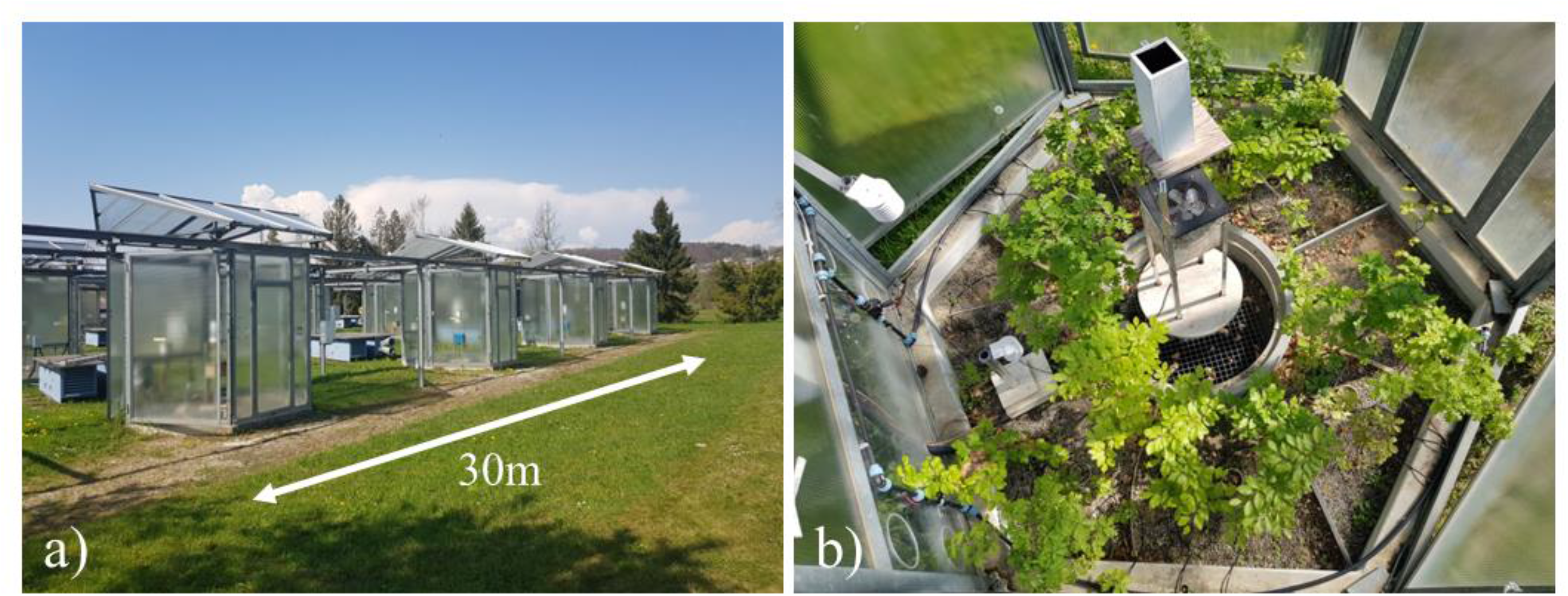
(a) Side picture of the 16 open-top chambers. (b) Aerial picture from a control chamber including a central fan and eight compartments with different species interaction treatments.

In October 2018, a total of 336 three-year-old seedlings of European beech and downy oak were planted into three species interaction treatments: single-trees to avoid any effects of species or individual interactions (n = 1 tree), monocultures to evaluate the effect of intraspecific interactions (i.e., four trees from the same species, n = 4 trees), and mixtures to assess the impact of interspecific interactions (i.e., diagonally two trees from each species, n = 4 trees). In this study, we only considered the monocultures and mixtures as we aimed to compare interaction effects. One tree per species was randomly selected for the measurement out of the two or four trees from the same species in each compartment. Tree seed originated from local nurseries from the canton of Solothurn (Biberist, Switzerland) and Valais (Leuk, Switzerland) for beech and oak trees, respectively, and were both grown by the compagny Schweizerpflanzen in the canton of Bern (Wiler bei Utzenstorf, Switzerland). Tree sizes were statistically similar when delivered and planted (Table **S4**). In June 2019, after the first leaf flushing and measurement campaign, trees were subjected to a combined manipulation of an air temperature and soil moisture regime. This resulted in four climatic treatments, including a control treatment with ambient air temperature and soil moisture maintained at field capacity (C), an air warming treatment with an increase of around 5°C (i.e., +5.19 °C ± 0.2 °C) relative to the (ambient) air temperature in the control (W) (leading also to an increase in VPD), a soil moisture reduction treatment with a reduction of soil moisture of around 50% (i.e., - 52.77 % ± 9.7 %) relative to the control (MR), and a combined air warming + soil moisture reduction treatment where both air warming and reduction of soil moisture were applied simultaneously (WMR, i.e., +5.02 °C ± 0.2 °C and - 51.96 % ± 10.3 %, Fig. **6**, Table **S5**). The soil field capacity was determined using pF-curves (corresponding to approx. 12% relative water content in this sandy soil). Our study aimed to understand the physiological mechanisms under chronic warming of +5°C and reduction of soil moisture conditions by 50% rather than predict the response of trees to periodic extreme events. The selected conditions have been chosen at our facility to match a possible future shift in mean air temperature leading also to constantly drier soils (Lyon et al., 2022). From March to November, trees were irrigated every second day using an automated irrigation system. Irrigation levels were adjusted throughout the year to maintain the soil moisture reduction in the low soil moisture and warming + soil moisture reduction treatments. To prevent frost damage to the pipes, the irrigation system was not used from December to March, and the watering was applied manually twice a month to maintain treatment differences. Air temperature and relative humidity were monitored inside each OTC at 50 cm and 2 m above the ground every 10 s, and averaged every 10 min (ATMOS 14, METER Group Inc., Pullman, WA, USA). Soil conditions, including soil temperature and moisture, were monitored at 25 cm depth every minute and averaged every 10 min (5TM, Decagon Devices, Pullman, WA, USA). Each of the four climatic treatments was repeated in four OTCs (n = 4). In each climatic treatment, beech and downy oak trees (n = 2 species) and species interaction treatments (monocultures and mixtures, n = 2) were replicated six times (n = 6 trees per climatic treatment, species and species interaction treatment = 96 trees in total). Phenology, leaf-level assimilation, starch and sugar concentrations, and growth were measured on the selected trees for three years (i.e., from 2019 to 2021). Phenology and growth were measured annually, and leaf-level parameters were measured three times a year during the growing season (i.e., early, middle, and late growing season). The first year of measurements (2019) was not included in this study because trees may still have been recovering from their transplant in the chambers. Moreover, phenological shifts were likely impacted by the climatic conditions of the previous years (e.g., Marchand et al., 2020), when the plants were still growing in the nursery.

### Phenology

From February to April, the timing of leaf flushing was monitored three times a week by the same observer. The bud development stages, from no bud activity to full leaf unfolding, were determined using a scale with five intermediate stages according to the species (Vitasse et al., 2009; Method **S1**). The bud swelling stage was reached in each tree when at least one bud reached stage 1. The development duration of leaf flushing represents the number of days needed to pass from stages 1 to 4. From September to December, the timing of leaf senescence was assessed once a week using the leaves’ coloration and leaf fall by the same observer. We considered that the onset of senescence was reached when trees had 10% of their leaves either colored or fallen, using a linear interpolation between two measuring dates, if necessary. The senescence duration was estimated as the number of days between the stage of 10% and 90% of either colored or fallen leaves. For each tree, the growing season length was calculated as the number of days between the bud swelling stage and the onset of senescence.

Growing degree-days (GDD) were calculated for each tree and year by accumulating the daily mean temperature above a threshold of 5°C from January 1^st^ to the date of bud swelling as determined in previous work on our focal tree species (Vitasse and Basler, 2013; Vitasse et al., 2019). The GDD requirement to budburst is an index used to determine the winter and spring temperatures that trigger budburst. GDD requirement to budburst is known to be influenced by the previous exposition of the plant to cooler temperatures, called chilling temperatures, that exponentially reduce the amount of GDD required to budburst (Murray et al., 1989). We also estimated the duration of chilling conditions for each tree during each winter by counting the number of days below a threshold of 5°C from November 1^st^ to the date of bud swelling.

### Foliar gas exchange

Measurements of light-saturated assimilation (A_sat_) were conducted by means of CO_2_ response curves (A/C_i_ curves) during the trees’ most active time of the day (i.e., between 9 am and 5 pm) using LI-COR 6800 infrared gas analyzers (LI-COR, Lincoln, USA), except in the late growing season of 2020, where LI-COR 6400 infrared gas analyzers were used. For the measurements, one fully mature and sun-exposed leaf per tree was selected from the top 1/3 of the crown. A_sat_ was extracted from the first point of the A/C_i_ curves at 400 ppm CO_2_ concentration, 1500 µmol m^-2^ s^-1^ PAR, block temperature matching mean daytime air temperature in the different treatments, and relative humidity at 50% (for the reference cell). The measurements are fully described in a previous study (Didion-Gency et al., 2022).

### Leaf water potential

Predawn and midday leaf water potential (Ψ_PD_, Ψ_MD_) were measured on one fully mature and sun-exposed leaf per tree from the top 1/3 of the crown that was stored in a plastic bag previously inhaled in. Predawn samples were collected before sunrise, and midday leaf water potential samples were collected in the middle of the day (solar time). Measurements were conducted on-site within 1.5 h after sample collection using a Scholander-type pressure chamber (PMS Instruments, USA).

### Non-structural carbohydrates

Leaves used for the midday leaf water potential measurements were microwaved at 600 W for 90 s and dried in the oven for at least 48 h at 65 °C until a stable weight was achieved. Leaves were then ground into a fine powder and used for non-structural carbohydrates (NSC) concentration measurements according to the previous explained protocols (Schönbeck et al., 2018; Method **S2**).

### Growth traits

Tree height and diameter were annually determined in September. Height was measured on the whole tree, and diameter was evaluated at the trunk base (15 cm above ground) using an electronic digital caliper. As no destructive measurements could be carried out in this ongoing experiment, we estimated the aboveground biomass (AGB), excluding leaves, with following the allometric equation from Annighöfer et al. (2016) (Method **S3**). The growth rate per year of each parameter was calculated by subtracting the current values to the value of the previous year. These parameters were then called height increment, diameter increment, and AGB increment.

### Biodiversity, complementarity and selection effects

Net biodiversity, complementarity, and selection effects were determined for each measured trait and species with following the equations of Loreau and Hector (2001) and Grossiord et al. (2013) (Method **S4**).

### Statistical analysis

The relationships between bud swelling, development duration, the onset of senescence, senescence duration, and growing season length, and the daily air temperature and soil moisture content were determined through linear regression (*lm* function) for both species in monocultures. Different air temperatures and soil moisture contents were tested to explain phenological variations: the daily mean air temperature and soil moisture content in spring (from February to April, T_spring_ and M_spring_, respectively), in fall (from September to December, T_fall_ and M_fall_, respectively), and annually (T_annual_ and M_annual_). Similarly, the relationship between the number of chilling days on the GDD requirement was determined through negative exponential regression (*nls* function) for both species in monocultures as this relationship is best represented by a negative exponential regression (Vitasse and Basler, 2013).

The response of all measured traits (i.e., starch concentration, sugar concentration, NSC concentration, A_sat_, height, diameter, and ABG increments) to the climatic treatments was determined through linear mixed-effects models for both species in monocultures. The interactive effects of warming (yes/no), soil moisture reduction (yes/no), and year (2019, 2020, and 2021) were used as fixed effects. As most leaf-level responses (i.e., assimilation, starch and sugar concentrations) did not vary during the growing season, we averaged them by years before the analyses. The individual chambers were treated as a random effect. Tukey-type posthoc tests were used to reveal significant differences between treatments for each year (*multcomp* function).

To determine the impact of species interactions on all our traits, a second linear mixed-effect model was conducted for both species using an additional species interaction effect (i.e., monocultures *vs*. mixtures) as a fixed effect. Similarly, Tukey-type posthoc tests were used to reveal significant differences between mixtures (*multcomp* function). When a significant species interaction effect was found, we used t-tests to determine if the net biodiversity, complementarity, and selection effects were significantly different from zero for each climatic treatment, year, and species (*t.test* function; Grossiord et al., 2013).

All analyses were performed using the R v.4.2.0 statistical platform (2022).

## Acknowledgements and funding

MD-G, CG, and YV were supported by the Swiss National Science Foundation SNF (PZ00P3_174068, 310030_204697, and 315230_192712). CG was further supported by the Sandoz Family Foundation.

## Author contributions

MD-G, and CG conceived and designed the study; MS and JG provided and managed the OTC facility and implemented measurement and control systems; MD-G and CG collected the data; MD-G analyzed the data and led the writing of the manuscript; All authors critically contributed to the manuscript and gave final approval for publication.

## Supplementary information

Supplementary information is available for this paper.

## Data availability statement

Data used in this manuscript will be available from the Dryad Digital Repository after acceptance (doi:10.5061/dryad.4j0zpc8hz). Data supporting the findings of this study are also available from the corresponding author, MD-G.

