## Supplementary information for "Chronic warming and dry soils limit carbon uptake and growth despite a longer growing season in beech and oak"

**Table S1:** Summary of the ANOVA tests of the linear mixed-effects models where the interactive effects of warming, moisture reduction, and year were evaluated on leaf-level carbon relations, and growth traits for beech and oak trees in monocultures.

**Table S2:** Summary of the ANOVA tests of the linear mixed-effects models where the interactive effects of warming, moisture reduction, year, and interaction were evaluated on phenological traits for beech and oak trees.

**Table S3:** Summary of the ANOVA tests of the linear mixed-effects models where the interactive effects of warming, moisture reduction, year, and interaction were evaluated on leaf-level carbon relations, and growth traits for beech and oak trees.

**Table S4:** Height and diameter for each species before trees grew in the different climatic.

**Table S5:** Summary of the climate for the different years, treatments, and times.

**Figure S1:** Phenological traits as a function of soil moisture for beech and oak trees growing under control, moisture reduction, warming, and warming + moisture reduction in the monocultures.

**Figure S2:** Selection effect on leaf-level carbon relations and growth traits for beech and oak trees growing under control, moisture reduction, warming, and warming + moisture reduction.

**Figure S3:** Complementarity effect on leaf-level carbon relations and growth traits for beech and oak trees growing under control, moisture reduction, warming, and warming + moisture reduction.

**Figure S4:** Description of the bud development for beech and oak trees.

**Supplementary Methods** **S1:** Description of bud development stages.

**Supplementary Methods** **S2**: Non-structural carbohydrates (NSC) concentration determination.

**Supplementary Methods** **S3:** Equation for the estimation of the aboveground biomass (AGB).

**Supplementary Methods S4:** Calculation of the net biodiversity, complementarity, and selection effect.

**Table S1:** Summary of the ANOVA tests of the linear mixed-effects models (F- and p-values) where the interactive effects of warming (W), moisture reduction (MR), and year (Y) were evaluated on the predawn leaf water potential (Ψ_PD_), midday leaf water potential (Ψ_MD_), starch, sugar, light-saturated assimilation (A_sat_), height increment, diameter increment, and estimated aboveground biomass (AGB) increment for beech and oak trees in monocultures. Significant effects (p ≤ 0.05) are highlighted in bold.

| Traits | Species | Sources of variations | | | | | | |
| --- | --- | --- | --- | --- | --- | --- | --- | --- |
|  |  | W | MR | Y | W * MR | W * Y | MR * Y | W * MR * Y |
| Ψ_PD_ | Beech | 2.83 (0.118) | 0.10 (0.761) | 0.25 (0.776) | 0.00 (0.983) | 2.12 (0.123) | 0.12 (0.891) | **3.54 (0.031)** |
|  | Oak | 0.34 (0.570) | 0.07 (0.797) | **7.95 (<0.001)** | 2.34 (0.152) | 0.90 (0.408) | 1.70 (0.185) | **3.59 (0.030)** |
| Ψ_MD_ | Beech | 0.96 (0.347) | 0.06 (0.804) | **6.50 (0.002)** | 0.04 (0.839) | 2.10 (0.123) | 0.27 (0.761) | 0.97 (0.381) |
|  | Oak | 0.50 (0.494) | 0.04 (0.839) | 2.25 (0.109) | 0.00 (0.984) | 0.35 (0.705) | 0.95 (0.387) | 0.42 (0.661) |
| Starch^*^ | Beech | 1.55 (0.236) | 0.44 (0.517) | 0.47 (0.623) | 0.10 (0.761) | 2.88 (0.059) | **3.48 (0.033)** | 2.14 (0.120) |
|  | Oak | 0.88 (0.366) | 0.69 (0.422) | 2.25 (0.109) | 0.57 (0.467) | 0.35 (0.708) | 0.12 (0.883) | 0.18 (0.839) |
| Sugar^*^ | Beech | 2.59 (0.133) | 1.34 (0.269) | 2.72 (0.069) | 2.04 (0.179) | 1.31 (0.273) | 1.89 (0.154) | 2.33 (0.100) |
|  | Oak | 2.55 (0.136) | 0.97 (0.343) | 3.15 (0.045) | 1.18 (0.298) | **3.70 (0.026)** | 0.56 (0.575) | 2.15 (0.119) |
| A_sat_^*^ | Beech | 2.44 (0.144) | 0.99 (0.339) | **25.86 (<0.001)** | 4.12 (0.065) | **5.22 (0.006)** | **19.00 (<0.001)** | 2.75 (0.067) |
|  | Oak | 0.00 (0.994) | 2.70 (0.127) | **3.41 (0.035)** | 0.17 (0.683) | 1.02 (0.363) | 0.66 (0.516) | 1.58 (0.209) |
| Height increment^*^ | Beech | 0.78 (0.395) | 0.50 (0.491) | 2.04 (0.133) | 0.03 (0.867) | 0.73 (0.483) | **13.82 (<0.001)** | **3.68 (0.027)** |
|  | Oak | 0.89 (0.363) | 1.08 (0.320) | **9.16 (<0.001)** | 1.23 (0.290) | **5.00 (0.008)** | **35.13 (<0.001)** | 1.90 (0.153) |
| Diameter increment^*^ | Beech | 1.45 (0.252) | 0.69 (0.422) | 2.24 (0.108) | 0.02 (0.898) | 0.62 (0.540) | **10.26 (<0.001)** | 0.85 (0.427) |
|  | Oak | **15.04 (0.002)** | **5.01 (0.045)** | **21.99 (<0.001)** | 2.85 (0.117) | **3.66 (0.026)** | **15.93 (<0.001)** | 1.88 (0.154) |
| AGB increment^*^ | Beech | 0.17 (0.690) | 0.04 (0.843) | 1.75 (0.185) | 0.09 (0.770) | 0.19 (0.826) | 2.83 (0.069) | 1.08 (0.348) |
|  | Oak | 0.33 (0.577) | 0.93 (0.355) | **5.05 (0.010)** | 0.89 (0.365) | 1.62 (0.209) | **7.27 (0.002)** | 1.04 (0.362) |
| Full model: *lme(Measured traits ~ (Warming + moisture reduction + Year)^3^, random = ~1\|Chamber)* | | | | | | | | |

^*^: square root transformation has been used.

**Table S2:** Summary of the ANOVA tests of the linear mixed-effects models (F- and p-values) where the interactive effects of warming (W), moisture reduction (MR), year (Y), and interaction (I; i.e., monocultures + mixtures) were evaluated on the bud swelling, leaf development duration, onset senescence, leaf senescence duration, and growing season length for beech and oak trees. Significant effects (p ≤ 0.05) are highlighted in bold.

| Traits | Species | Sources of variations | | | | | | |
| --- | --- | --- | --- | --- | --- | --- | --- | --- |
|  |  | I | W * I | MR * I | Y * I | W * MR * I | W * Y * I | MR * Y * I |
| Bud swelling^*^ | Beech | 2.11 (0.150) | 2.71 (0.104) | 0.62 (0.432) | 0.02 (0.898) | 0.80 (0.373) | 0.04 (0.849) | 0.18 (0.676) |
|  | Oak | 0.31 (0.581) | 1.14 (0.289) | 0.36 (0.550) | 0.01 (0.907) | 1.48 (0.228) | 1.10 (0.298) | 0.01 (0.931) |
| Development duration^*^ | Beech | 0.28 (0.598) | 2.18 (0.144) | 0.05 (0.815) | 0.81 (0.372) | 0.00 (0.994) | 1.19 (0.279) | 0.76 (0.388) |
|  | Oak | 0.67 (0.415) | 0.20 (0.655) | 0.09 (0.765) | **9.37 (0.003)** | 3.46 (0.067) | 1.09 (0.300) | **11.21 (0.001)** |
| Onset senescence^*^ | Beech | 0.11 (0.736) | 0.01 (0.927) | 1.10 (0.298) | 0.18 (0.837) | 0.78 (0.378) | 0.29 (0.752) | 0.89 (0.414) |
|  | Oak | 0.71 (0.401) | 0.00 (0.948) | 1.65 (0.202) | 0.27 (0.763) | 1.18 (0.280) | 2.07 (0.131) | 1.59 (0.209) |
| Senescence duration^*^ | Beech | 1.44 (0.232) | 0.15 (0.695) | 0.00 (0.983) | 0.16 (0.849) | 0.48 (0.491) | 0.18 (0.835) | 0.10 (0.908) |
|  | Oak | 0.17 (0.680) | 0.46 (0.497) | 2.88 (0.092) | 0.83 (0.439) | 1.35 (0.249) | 0.66 (0.520) | 0.35 (0.706) |
| Growing season length^*^ | Beech | 0.52 (0.473) | 0.49 (0.488) | 0.96 (0.330) | 0.37 (0.694) | 2.18 (0.144) | 0.79 (0.455) | 0.13 (0.877) |
|  | Oak | 1.87 (0.174) | 0.33 (0.564) | 0.29 (0.593) | 2.24 (0.112) | 0.55 (0.460) | 0.04 (0.963) | 0.54 (0.586) |
| Full model: *lme(Measured traits ~ (Warming + moisture reduction + Year + Interaction)^3^, random = ~1\|Chamber)* | | | | | | | | |

^*^: square root transformation has been used.

**Table S3:** Summary of the ANOVA tests of the linear mixed-effects models (F- and p-values) where the interactive effects of warming (W), moisture reduction (MR), year (Y) and interaction (I; i.e., monocultures + mixtures) were evaluated on the starch, sugar, light-saturated assimilation (A_sat_), height increment, diameter increment, and estimated aboveground biomass (AGB) increment for beech and oak trees. Significant effects (p ≤ 0.05) are highlighted in bold.

| Traits | Species | Sources of variations | | | | | | |
| --- | --- | --- | --- | --- | --- | --- | --- | --- |
|  |  | I | W * I | MR * I | Y * I | W * MR * I | W * Y * I | MR * Y * I |
| Starch^*^ | Beech | 0.32 (0.572) | 0.34 (0.558) | 2.35 (0.126) | 0.42 (0.659) | 0.04 (0.838) | 0.20 (0.822) | 2.34 (0.098) |
|  | Oak | 0.00 (0.963) | 0.02 (0.888) | 0.63 (0.428) | 0.38 (0.683) | 1.08 (0.300) | 0.24 (0.790) | 0.08 (0.921) |
| Sugar^*^ | Beech | 0.12 (0.730) | 0.08 (0.774) | 0.12 (0.730) | 2.37 (0.095) | 0.00 (0.994) | 1.16 (0.315) | 0.30 (0.744) |
|  | Oak | 0.63 (0.427) | 1.21 (0.271) | 0.59 (0.444) | 0.15 (0.862) | 0.25 (0.616) | 1.03 (0.357) | 0.05 (0.951) |
| A_sat_^*^ | Beech | 1.03 (0.310) | 2.34 (0.127) | 2.02 (0.156) | 1.50 (0.224) | 3.65 (0.057) | 1.13 (0.323) | 1.94 (0.144) |
|  | Oak | 0.60 (0.438) | 0.55 (0.459) | 0.29 (0.590) | 0.02 (0.977) | 0.02 (0.878) | 0.97 (0.378) | 1.47 (0.231) |
| Height increment^*^ | Beech | 0.26 (0.611) | **6.80 (0.009)** | 0.68 (0.411) | 2.41 (0.092) | **11.91 (0.001)** | 1.19 (0.307) | 2.42 (0.090) |
|  | Oak | 0.00 (0.975) | 1.53 (0.217) | 0.28 (0.599) | 0.64 (0.530) | 0.66 (0.417) | 0.09 (0.916) | 0.29 (0.748) |
| Diameter increment^*^ | Beech | 0.27 (0.603) | 0.00 (0.968) | 3.66 (0.057) | **3.08 (0.047)** | 0.19 (0.665) | 0.52 (0.596) | **15.06 (<0.001)** |
|  | Oak | 2.11 (0.148) | 0.84 (0.360) | 0.08 (0.776) | 1.75 (0.175) | 0.99 (0.320) | **4.15 (0.016)** | 1.42 (0.243) |
| AGB increment^*^ | Beech | 0.02 (0.897) | 2.61 (0.109) | 0.06 (0.801) | 0.56 (0.574) | 3.60 (0.060) | 0.80 (0.454) | 0.69 (0.504) |
|  | Oak | 0.01 (0.940) | 0.11 (0.739) | 0.04 (0.841) | 1.60 (0.206) | 0.02 (0.878) | 0.20 (0.820) | 1.73 (0.181) |
| Full model: *lme(Measured traits ~ (Warming + moisture reduction + Year + Interaction)^3^, random = ~1\|Chamber)* | | | | | | | | |

^*^: square root transformation has been used.

**Table S4:** Average and standard deviation of the height and diameter for each species in spring 2019 (i.e., before the start of the climatic treatments) in control (C), moisture reduction (MR), warming (W), and warming + moisture reduction (WMR) treatments.

| **Species** | **Treatment** | **Height (cm)** | **Diameter (mm)** |
| --- | --- | --- | --- |
| Beech | C | 66.7 ± 13.2 | 6.0 ± 1.6 |
|  | MR | 66.3 ± 10.8 | 6.1 ± 1.3 |
|  | W | 64.9 ± 7.4 | 5.2 ± 1.3 |
|  | WMR | 61.1 ± 11.6 | 5.5 ± 1.4 |
| Oak | C | 66.6 ± 11.4 | 5.7 ± 1.5 |
|  | MR | 65.3 ± 14.7 | 5.5 ± 1.5 |
|  | W | 64.3 ± 13.2 | 5.4 ± 1.5 |
|  | WMR | 68.9 ± 12.9 | 5.8 ± 1.3 |

**Table S5:** Average and standard deviation of the air temperature, vapor pressure deficit (VPD), and soil moisture content under control (C), moisture reduction (MR), warming (W), and warming + moisture reduction (WMR) treatments measured in the OTCs each year in spring (from February to April), autumn (from September to December), annually (from January to December), and during the measurement campaigns (C1, C2, etc.).

| **Year** |  | **Air temperature (°C)** | | | |  | **VPD (kPa)** | | | |  | **Soil moisture (%)** | | | |
| --- | --- | --- | --- | --- | --- | --- | --- | --- | --- | --- | --- | --- | --- | --- | --- |
|  |  | C | MR | W | WMR |  | C | MR | W | WMR |  | C | MR | W | WMR |
| 2019 | Annual | 11.2 ± 0.4 | 11.3 ± 0.4 | 14.3 ± 0.5 | 14.2 ± 0.5 |  | 0.4 ± 0.0 | 0.4 ± 0.0 | 0.9 ± 0.0 | 0.9 ± 0.0 |  | 9.3 ± 0.1 | 6.5 ± 0.0 | 9.7 ± 0.1 | 6.5 ± 0.0 |
|  | Spring | 6.9 ± 0.5 | 6.9 ± 0.5 | 7.0 ± 0.5 | 7.0 ± 0.5 |  | 0.6 ± 0.1 | 0.6 ± 0.1 | 0.6 ± 0.1 | 0.6 ± 0.1 |  | 7.8 ± 0.0 | 8.1 ± 0.1 | 6.6 ± 0.1 | 7.5 ± 0.0 |
|  | Autumn | 9.5 ± 0.5 | 9.7 ± 0.5 | 14.6 ± 0.5 | 14.5 ± 0.5 |  | 0.2 ± 0.0 | 0.2 ± 0.0 | 0.6 ± 0.0 | 0.6 ± 0.0 |  | 10.2 ± 0.1 | 6.1 ± 0.0 | 10.5 ± 0.1 | 6.3 ± 0.0 |
|  | C1 | 14.0 ± 4.3 | 14.4 ± 4.6 | 14.6 ± 4.7 | 14.3 ± 4.5 |  | 0.3 ± 0.2 | 0.4 ± 0.3 | 0.4 ± 0.3 | 0.4 ± 0.2 |  | 6.9 ± 0.8 | 7.1 ± 0.9 | 7.4 ± 0.9 | 6.6 ± 0.8 |
|  | C2 | 19.4 ± 5.0 | 19.7 ± 5.2 | 24.7 ± 5.2 | 24.4 ± 5.1 |  | 0.7 ± 0.2 | 0.7 ± 0.2 | 1.4 ± 0.3 | 1.3 ± 0.3 |  | 9.3 ± 2.4 | 6.2 ± 0.9 | 10.3 ± 2.8 | 6.4 ± 1.8 |
|  | C3 | 17.2 ± 6.1 | 17.5 ± 6.2 | 22.4 ± 6.2 | 22.2 ± 6.1 |  | 0.5 ± 0.1 | 0.5 ± 0.1 | 1.1 ± 0.1 | 1.1 ± 0.1 |  | 10.5 ± 3.0 | 6.1 ± 0.2 | 10.8 ± 2.0 | 6.5 ± 2.2 |
| 2020 | Annual | 11.8 ± 0.4 | 12.1 ± 0.4 | 17.0 ± 0.4 | 16.8 ± 0.4 |  | 0.4 ± 0.0 | 0.4 ± 0.0 | 0.9 ± 0.0 | 0.9 ± 0.0 |  | 10.3 ± 0.0 | 5.3 ± 0.0 | 10.3 ± 0.0 | 4.8 ± 0.0 |
|  | Spring | 9.0 ± 0.5 | 9.3 ± 0.5 | 14.2 ± 0.5 | 14.1 ± 0.5 |  | 0.4 ± 0.0 | 0.4 ± 0.0 | 0.8 ± 0.0 | 0.8 ± 0.0 |  | 9.8 ± 0.1 | 5.3 ± 0.0 | 9.9 ± 0.1 | 4.9 ± 0.0 |
|  | Autumn | 8.9 ± 0.6 | 9.1 ± 0.6 | 14.1 ± 0.6 | 13.9 ± 0.6 |  | 0.2 ± 0.0 | 0.2 ± 0.0 | 0.6 ± 0.0 | 0.6 ± 0.0 |  | 10.8 ± 0.0 | 5.4 ± 0.0 | 10.7 ± 0.0 | 5.0 ± 0.0 |
|  | C4 | 16.5 ± 3.6 | 16.8 ± 3.9 | 21.7 ± 3.9 | 21.4 ± 3.6 |  | 0.4 ± 0.2 | 0.4 ± 0.2 | 0.9 ± 0.2 | 0.9 ± 0.2 |  | 10.2 ± 2.3 | 4.9 ± 0.9 | 10.6 ± 2.2 | 4.3 ± 0.7 |
|  | C5 | 23.7 ± 5.4 | 24.1 ± 5.5 | 28.9 ± 5.5 | 28.6 ± 5.5 |  | 1.0 ± 0.1 | 1.0 ± 0.1 | 1.8 ± 0.2 | 1.7 ± 0.2 |  | 10.4 ± 2.0 | 5.2 ± 1.1 | 10.4 ± 2.3 | 4.7 ± 1.2 |
|  | C6 | 19.6 ± 6.0 | 19.9 ± 6.1 | 24.7 ± 6.0 | 24.6 ± 6.0 |  | 0.5 ± 0.1 | 0.6 ± 0.1 | 1.2 ± 0.1 | 1.2 ± 0.1 |  | 10.8 ± 2.0 | 5.2 ± 1.5 | 10.8 ± 2.4 | 4.9 ± 1.5 |
| 2021 | Annual | 10.5 ± 0.4 | 10.8 ± 0.4 | 15.7 ± 0.4 | 15.6 ± 0.4 |  | 0.3 ± 0.0 | 0.3 ± 0.0 | 0.8 ± 0.0 | 0.8 ± 0.0 |  | 10.0 ± 0.0 | 4.6 ± 0.0 | 10.3 ± 0.0 | 4.8 ± 0.0 |
|  | Spring | 6.7 ± 0.5 | 6.9 ± 0.5 | 11.9 ± 0.5 | 11.7 ± 0.5 |  | 0.3 ± 0.0 | 0.3 ± 0.0 | 0.7 ± 0.0 | 0.7 ± 0.0 |  | 9.7 ± 0.1 | 5.0 ± 0.0 | 10.3 ± 0.1 | 4.1 ± 0.0 |
|  | Autumn | 8.7 ± 0.6 | 8.9 ± 0.6 | 13.9 ± 0.6 | 13.8 ± 0.6 |  | 0.2 ± 0.0 | 0.2 ± 0.0 | 0.6 ± 0.0 | 0.6 ± 0.0 |  | 10.1 ± 0.0 | 4.3 ± 0.0 | 10.4 ± 0.0 | 4.9 ± 0.1 |
|  | C7 | 22.8 ± 6.4 | 23.3 ± 6.6 | 28.1 ± 6.5 | 27.6 ± 6.5 |  | 0.9 ± 0.2 | 1.0 ± 0.2 | 1.7 ± 0.2 | 1.7 ± 0.2 |  | 9.8 ± 1.6 | 4.8 ± 1.4 | 9.8 ± 2.0 | 5.4 ± 1.6 |
|  | C8 | 19.4 ± 3.6 | 19.7 ± 3.7 | 24.5 ± 3.7 | 24.4 ± 3.6 |  | 0.4 ± 0.1 | 0.4 ± 0.1 | 1.0 ± 0.2 | 1.0 ± 0.2 |  | 10.0 ± 1.7 | 4.6 ± 1.3 | 10.0 ± 2.1 | 5.3 ± 1.7 |
|  | C9 | 18.7 ± 5.7 | 19.0 ± 6.0 | 23.8 ± 5.9 | 23.7 ± 6.1 |  | 0.6 ± 0.1 | 0.6 ± 0.1 | 1.1 ± 0.1 | 1.2 ± 0.1 |  | 10.5 ± 1.8 | 4.5 ± 1.1 | 10.6 ± 2.1 | 5.7 ± 2.3 |

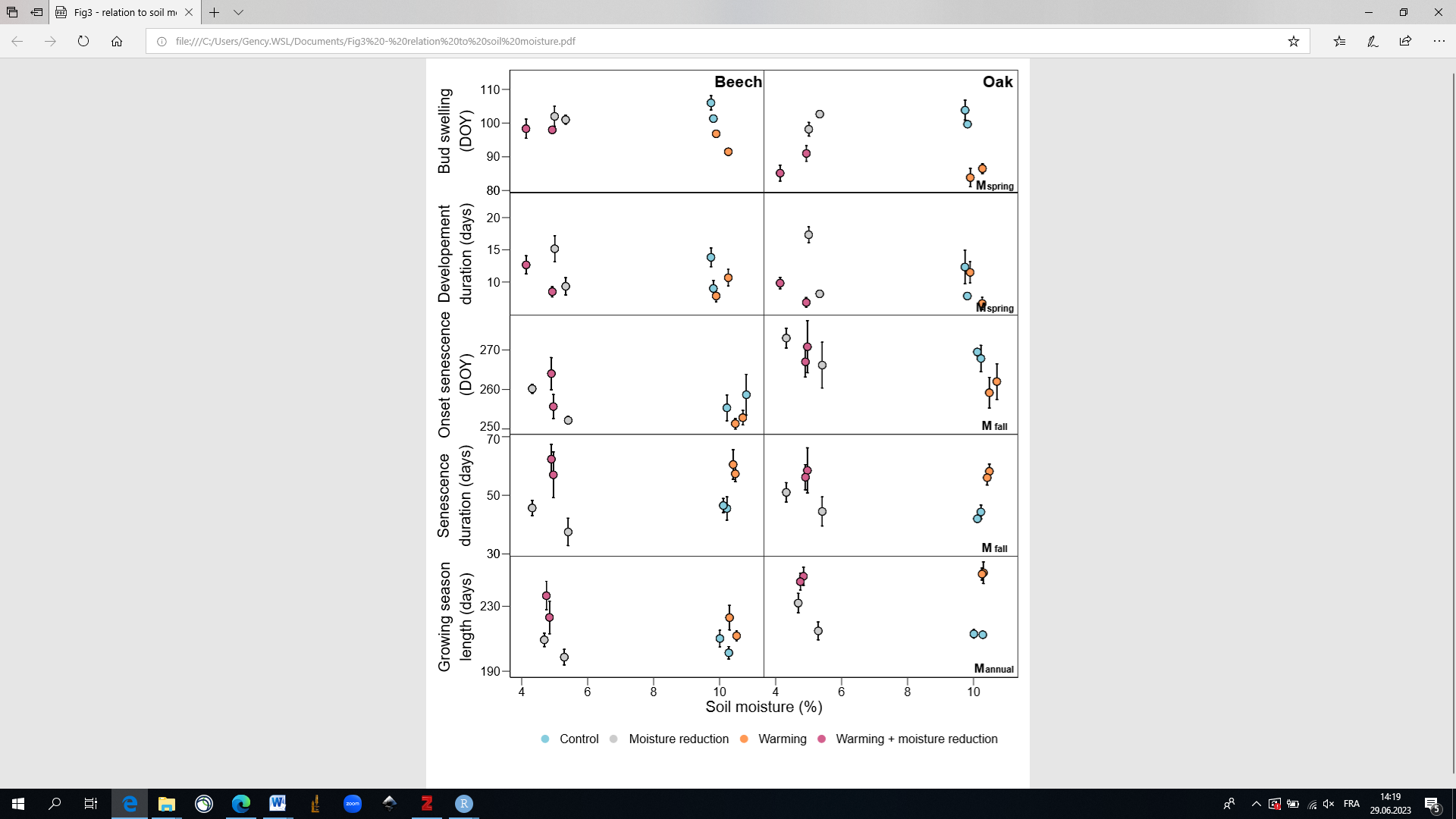

**Figure S1:** Bud swelling, leaf development duration, onset of senescence, leaf senescence duration, and growing season length as a function of soil moisture for beech and oak trees growing under control (blue), moisture reduction (grey), warming (orange), and warming + moisture reduction (purple) in the monocultures (mean ± SE per treatment and species for the years 2020 and 2021, n = 6 trees). The bud swelling and leaf development duration are shown as a function of the mean soil moisture measured in early spring (from February to April, M_spring_). Onset of senescence and leaf senescence duration are shown as a function of the mean soil moisture in fall (from September to December, M_fall_). Growing season length is shown as a function of the mean annual soil moisture (M_annual_). Linear regression lines across all treatments per species are shown when significant. R², p-values and the change of day number for each degree are given when significant.

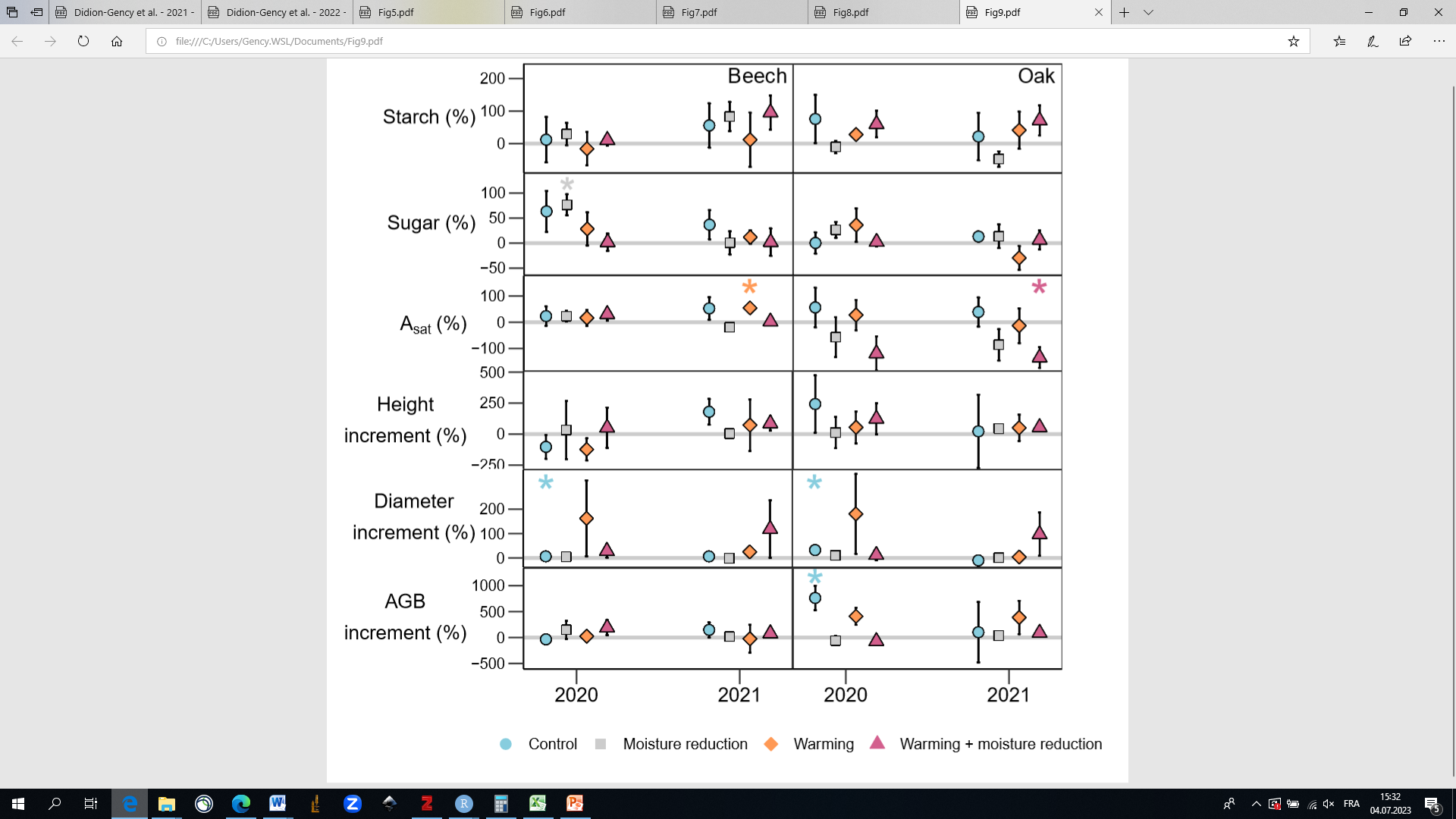

**Figure S2:** Starch, sugar, light‐saturated assimilation (A_sat_), height increment, diameter increment, and estimated aboveground biomass (AGB) increment selection effect for beech and oak trees growing under control (blue), moisture reduction (grey), warming (orange), and warming + moisture reduction (purple) (mean ± SE per treatment and species for the years 2020 and 2021, n = 18 trees for the physiological traits and, n = 6 trees for the growth traits). Positive values indicate higher rates in mixtures compared to the monocultures. Only significant differences from 0 are highlighted per treatment, and species (p - value: 0.05 ≥ * > 0.01, 0.01 ≥ ** > 0.001, *** ≥ 0.001).

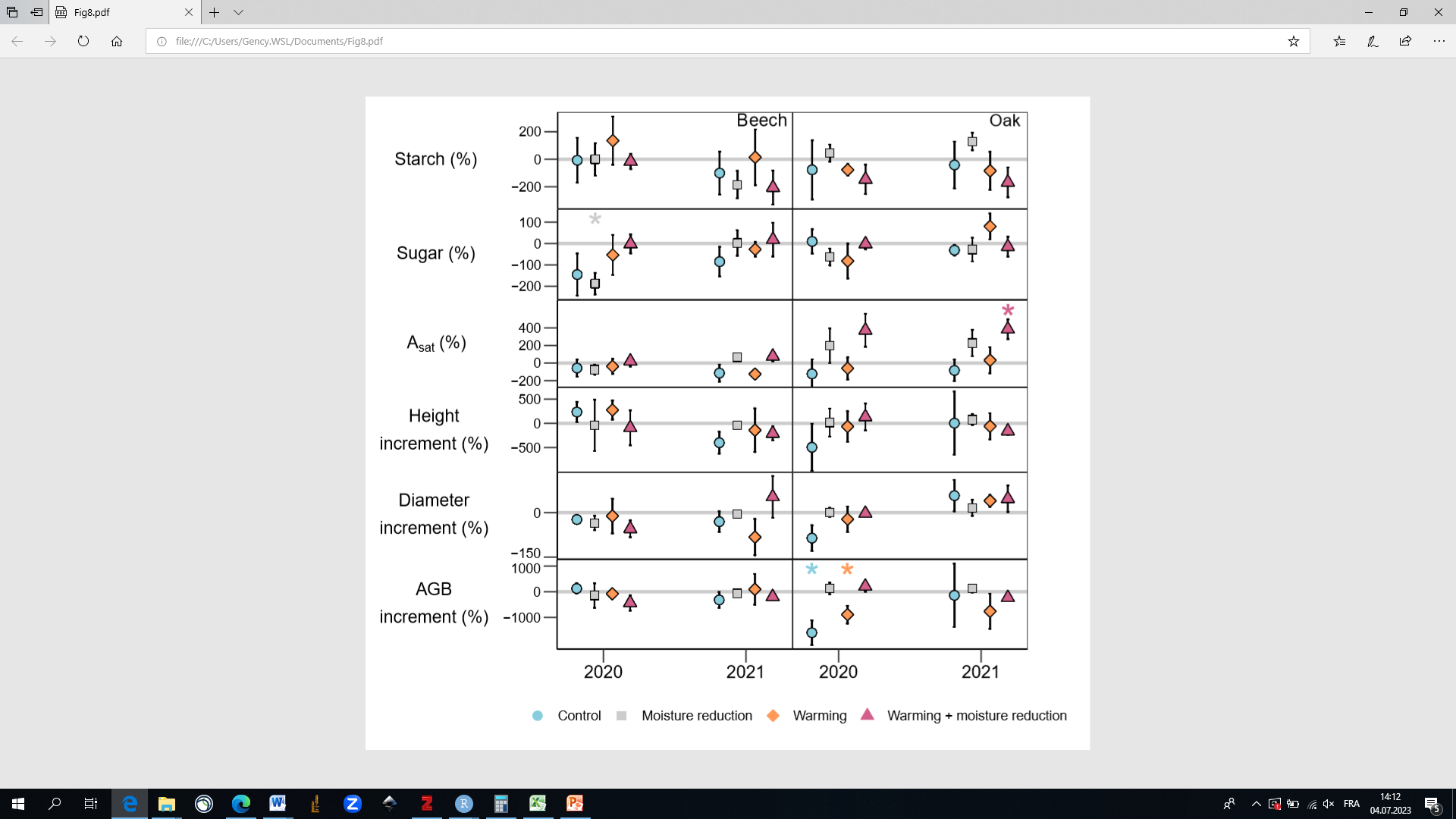

**Figure S3:** Starch, sugar, light‐saturated assimilation (A_sat_), height increment, diameter increment, and estimated aboveground biomass (AGB) increment complementarity effect for beech and oak trees growing under control (blue), moisture reduction (grey), warming (orange), and warming + moisture reduction (purple) (mean ± SE per treatment and species for the years 2021, n = 18 trees for the physiological traits and n = 6 trees for the growth traits). Positive values indicate higher rates in mixtures compared to the monocultures. Only significant differences from 0 are highlighted per treatment, and species (p - value: 0.05 ≥ * > 0.01, 0.01 ≥ ** > 0.001, *** ≥ 0.001).

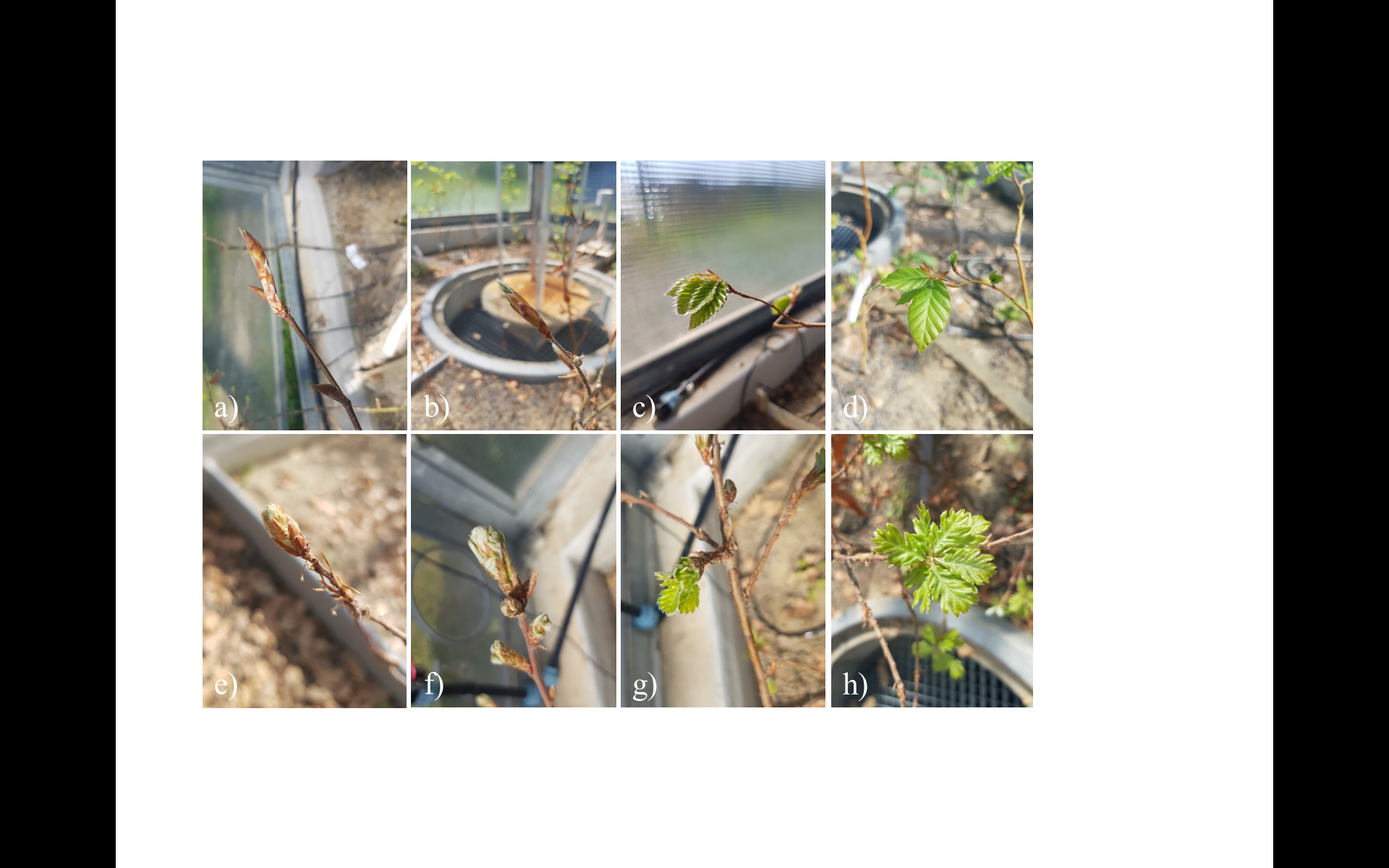

**Figure S4:** Bud development at stage 1 (a, e), stage 2 (b, f), stage 3 (c, g), and stage 4 (d, h) for beech and oak trees, respectively.

**Supplementary Methods** **S1:**

**Description of bud development stages**

At stage 1, buds became softer and greenish; at stage 2, tips of the leaf came out; at stage 3, the green of the new leaves is visible; and at stage 4, leaves are fully visible and at least one is completely unfolded (Fig. **S4**).

**Supplementary Methods** **S2:**

**Non-structural carbohydrates (NSC) concentration determination.**

Ten to twelve mg of ground leaf material were steamed in 2 mL distilled water for 30 min. An aliquot of 200 µL was treated for an hour with Invertase from baker's yeast (*S. cerevisiae*, Sigma-Aldrich Chemie GmbH, Germany) to degrade sucrose and convert fructose into glucose. The sugar concentration was determined at 340 nm in a 96-well plate spectrophotometer (Thermo Fisher Scientific Multiskan GO, Finland) after an enzymatic conversion to gluconate-6-phosphate using Isomerase from baker's yeast (S. cerevisiae, Sigma-Aldrich Chemie GmbH, Germany) and glucose Assay Reagent (Sigma-Aldrich Chemie GmbH, Germany). The total amount of NSC was measured by taking an aliquot of 500 µL of the extract (including starch and sugar) and treated for 15 h at 49 °C with Amyloglucosidase from *Aspergillus niger* (Sigma-Aldrich Chemie GmbH, Germany) to digest starch into glucose. Total NSC concentration was determined using a spectrophotometer, as explained above. The starch concentration was calculated as the total NSC subtracted by the sugar concentration. Standard solutions, including pure starch, glucose, fructose, sucrose, and standard plant powder (Orchard leaves; Leco, USA), were used to allow the comparison and reproducibility between runs.

**Supplementary Methods** **S3:**

**Equation for the estimation of the aboveground biomass (AGB).**

$AGB= \beta1 H^{\beta2}$	Eq. 1

where *H* is the tree height, and *β1* and *β2* are species-specific coefficients for beech and oak trees (Annighöfer et al., 2016). The tree height, diameter, and AGB increments were calculated by subtracting the current year from the previous year’s measurements.

**Supplementary Methods** **S4:**

**Calculation of the net biodiversity, complementarity, and selection effect.**

$Net biodiversity effect (\%)= F_{O}- F_{E}={[(F}_{O}* W_{O})-{(F}_{E}* W_{O})]* 100$	Eq. 2

and

$Complementarity effect (\%)= [N *\left( \frac{F_{O}* W_{O}}{F_{E}}-W_{O} \right)* F_{E}]* 100$	Eq. 3

and

$Selection effect (\%)=[\left( \frac{F_{O}* W_{O}}{F_{E}}-W_{O} \right)- \left( \frac{F_{O}* W_{O}}{F_{E}}-W_{O} \right)* F_{E}]* 100$	Eq. 4

where F_O_ is the observed trait value of the species *i* in mixtures, *W_O_* is the proportion of species *i* in the mixtures in terms of the number of individuals (i.e., 0.5), *F_E_* is the observed trait value of species *i* in the monocultures, and *N* is the number of species.
